# Novel taxa of Acidobacteriota involved in seafloor sulfur cycling

**DOI:** 10.1101/2020.10.01.322446

**Authors:** Mathias Flieder, Joy Buongiorno, Craig W. Herbold, Bela Hausmann, Thomas Rattei, Karen G. Lloyd, Alexander Loy, Kenneth Wasmund

## Abstract

Acidobacteriota are widespread and often abundant in marine sediments, yet their metabolic and ecological properties are poorly understood. Here, we examined metabolisms and distributions of Acidobacteriota in marine sediments of Svalbard by functional predictions from metagenome-assembled genomes (MAGs), amplicon sequencing of 16S rRNA and dissimilatory sulfite reductase (*dsrB*) genes and transcripts, and gene expression analyses of tetrathionate-amended microcosms. Acidobacteriota were the second most abundant *dsrB*-harboring (averaging 13%) phylum after Desulfobacterota in Svalbard sediments, and represented 4% of *dsrB* transcripts on average. We propose two new Acidobacteriota genera, *Candidatus* Sulfomarinibacter (class Thermoanaerobaculia, ‘sub-division 23’) and *Ca*. Polarisedimenticola (‘sub-division 22’), with distinct genetic properties that may explain their distributions in biogeochemically distinct fjord sediments. *Ca*. Sulfomarinibacter encodes flexible respiratory routes, with potential for oxygen, nitrous oxide, metal-oxide, tetrathionate, sulfur and sulfite/sulfate respiration, and possibly sulfur disproportionation. Potential nutrients and energy include cellulose, proteins, cyanophycin, hydrogen and acetate. A *Ca*. Polarisedimenticola MAG encodes enzymes to degrade proteins, and to reduce oxygen, nitrate, sulfur/polysulfide and metal-oxides. 16S rRNA gene and transcript profiling showed *Ca*. Sulfomarinibacter members were relatively abundant and transcriptionally active in sulfidic fjord sediments, while *Ca*. Polarisedimenticola members were more relatively abundant in metal-rich fjord sediments. Overall, we reveal various physiological features of uncultured marine Acidobacteriota that indicate fundamental roles in seafloor biogeochemical cycling.

## Introduction

Bacteria of the phylum *Acidobacteriota* (also known as ‘Acidobacteria’) are highly diverse and inhabit a vast array of environments on Earth, yet the properties of various Acidobacteriota lineages remain poorly understood [1–6]. Knowledge regarding the functions and ecology of Acidobacteriota is biased to isolates and genomes obtained from soils, where they are especially prevalent and often dominate microbial communities [3, 4]. Soil-derived Acidobacteriota are generally known as aerobic heterotrophs that utilize various carbohydrates including polysaccharides like chitin or cellulose [3, 7, 8]. Some Acidobacteriota known from other environments have unique physiological properties, such as the ability to reduce iron [9], perform phototrophy [9, 10], or exhibit thermophilic lifestyles [11]. Members of Acidobacteriota sub-divisions 1 and 3 from peatland and permafrost soils have the potential to dissimilate inorganic and/or organic sulfur compounds [2, 12]. In comparison to terrestrial Acidobacteriota, even less is known about Acidobacteriota in marine systems.

Acidobacteriota 16S rRNA genes or genomes are frequently detected in marine environments including ocean waters, marine sponges, hydrothermal vents, or sediments [13–17]. Studies of 16S rRNA genes in marine sediments showed that Acidobacteriota are widespread and reach relative abundances in amplicon libraries of up to 23% [18–23]. This suggests they play important roles in microbial community functioning and biogeochemical processes, although our knowledge regarding their specific roles in sediments remains limited. A recent stable isotope probing study showed some Acidobacteriota in deep-sea sediments are capable of fixing nitrogen [24]. Acidobacteriota were also shown to be active, by incorporation of isotopically-labelled tracer into their DNA, under sulfidic conditions in incubations with estuarine sediment [6]. One novel Acidobacteriota metagenome-assembled genome (MAG) (*Candidatus* Guanabacteria) had genes for the CO dehydrogenase/CO-methylating acetyl-CoA synthase complex and heterodisulfide reductases, indicating a possible anaerobic lifestyle [25].

Marine sediments are a massive global habitat for microorganisms [26], with cell densities of microorganisms average up to 10^9^ cells per cm^3^ in surface sediments of organic-rich sediments [27]. Substantial amounts of organic matter are processed in marine sediments, which makes them a critical component of marine and global biogeochemical cycles [28]. Marine sediments are often stratified with respect to redox states, whereby oxygen is typically depleted within millimetres to centimetres below the surface at sites where organic inputs are relatively high [29]. Vast expanses of sediments are therefore anoxic, and many microorganisms survive via anaerobic lifestyles, such as fermentation, or respiration of nitrate, metals, sulfate or CO_2_. Sulfate is abundant in sediments and is used by sulfite/sulfate-reducing microorganisms (SRMs) as an electron acceptor for anaerobic respiration. Sulfate reduction is estimated to facilitate approximately 29% of organic matter degradation in marine sediments globally [26, 28]. The sulfur cycle is therefore a major driver of microbial life and biogeochemical cycling in the seafloor, so understanding the microorganisms that catalyze sulfur cycling is of great importance.

Because sulfate reduction is a major process in marine sediments, the activities, distributions and diversity of SRMs have been relatively well studied [28, 30, 31]. Members of the Desulfobacterota (formerly ‘Deltaproteobacteria’) are known as abundant SRMs in marine sediments, playing key roles in anaerobic food webs by utilizing fermentation products released by primary degraders of organic matter [32–34]. They are also represented by various isolates, and many have been subject to genomic and physiological studies [35]. Surveys of functional marker genes for sulfite/sulfate reducers in marine sediments, i.e., of dissimilatory sulfite reductases (*dsrAB*), have repeatedly shown that *dsrAB* from the phylum Desulfobacterota are typically the dominant *dsrAB*-harbouring group in marine sediments, but importantly, that several other lineages of *dsrAB*-harbouring uncultivated organisms are also abundant and prevalent [36]. Recently, some *dsrAB* sequences in marine sediments have been inferred to belong to Acidobacteriota [6, 37], although nothing is known about the metabolic properties or the sulfur dissimilating pathways of the organisms that harbour these genes. Identifying and understanding these undescribed *dsrAB*-harbouring microorganisms is therefore critical for understanding the microbial groups that drive sulfur cycling in marine sediments.

In this study, we aimed to gain insights into the metabolic potential of uncultured Acidobacteriota in marine sediments. We therefore recovered metagenome-assembled genomes (MAGs) from abundant Acidobacteriota populations present in marine fjord sediments of Svalbard, and predicted their metabolic features. Focus was placed on MAGs from the Thermoanaerobaculia, which represent a newly described lineage of *dsrAB*-harbouring organisms that may be important sulfur cycling bacteria in marine sediments. These analyses were complemented with comparative genomics, incubation experiments, transcript analysis, and analyses of Acidobacteriota distributions in Svalbard sediments, together revealing they may play various roles in sedimentary biogeochemical cycles, and that they are a prominent group of sulfur-dissimilating organisms.

## Materials and Methods

### Sample collection and microcosms

Marine sediments were collected from Smeerenburgfjorden, Kongsfjorden and Van Keulenfjorden, Svalbard, Norway, in July 2016 and/or June 2017 with the vessel ‘MS Farm’. Extensive biogeochemical data for these sites is available from previous studies [38–42]. Maps of sample locations are presented in Michaud *et al*. 2020. From Smeerenburgfjorden, samples were taken from three stations: station GK (79°38.49N, 11°20.96E), station J (79°42.83N, 11°05.10E) and station GN (79°45.01N, 11°05.99E). Samples from Van Keulenfjorden were taken from sites AC (77°32.260’N, 15°39.434’E) and AB (77°35.249’N, 15°05.121’E). A sample was also taken from Kongsfjorden station F (78°55.075’ N, 12°15.929’ E) [43]. For molecular biological analyses, samples were taken with HAPS [44] or Rumohr corers [45]. Details of core subsampling procedures and microcosm incubations are provided in the Supplementary information. Additional samples for non-quantitative microscopy were taken from tidal flat sediments of Aveiro Lagoon, Portugal (40°34.14N 8°45.10W) in July 2019, and from Kristineberg station near Fiskebäckskil, Sweden (58°24.95N, 11°44.50E) in October 2019, with further details provided in the Supplementary information.

### Nucleic acid extractions and reverse-transcription

For amplicon-based analyses, DNA and RNA was extracted from the sediment core samples (*∼*500 µl) and microcosm samples (*∼*250 µl) using the RNeasy PowerSoil Total RNA Kit (Qiagen) according to the manufacturer’s instructions. Additionally, a phenol/chloroform based extraction method was used to extract nucleic acids from sediment samples from station J sampled in July 2016 (Supplementary information). Eluted nucleic acids were stored in molecular biology grade water at –80°C. Aliquots for DNA-based analyses were used as eluted, while aliquots for RNA-based analyses were DNase-treated using the TURBO DNA-free™ kit (Thermo Fisher), followed by reverse transcription of the RNA to cDNA using the RevertAid First Strand cDNA Synthesis Kit (Thermo Fisher) according to the manufacturer’s instructions. To test if any DNA remained in the RNA samples after the DNase digestion step, control samples were processed as above except the RevertAid M-MuLV Reverse Transcriptase was excluded. These controls were checked for DNA by PCR using 16S rRNA gene targeting primers (described below).

Sediment samples from 2016 were used for metagenome sequencing. DNA was extracted by the Vienna group from 3–5 mL of sediment from varying depths or microcosms derived from station J, Smeerenburgfjorden, and 18 centimeters below seafloor (cmbsf) from station AC of Van Keulenfjorden (Supp. Table 1) using the DNeasy PowerSoil Kit (Qiagen) according to the manufacturer’s protocol. DNA was also extracted by the Knoxville group from 2 grams of sample spanning 0–5 cmbsf from site AB of Van Keulenfjorden and site F of Kongsfjorden (Supp. Table 1), using the RNeasy PowerSoil Kit (Qiagen) with DNA elution following the manufacturer’s protocol.

### Metagenome sequencing and genome binning

DNA libraries were prepared (detailed in Supplementary information) and sequenced using 2×150 bp paired-end mode on an Illumina HiSeq 3000 instrument at the Biomedical Sequencing Facility (BSF), Vienna. Metagenomic libraries were generated from the combined extracts from the first 5 cm (spanning 0 to 5 cm downcore) in sites AB and F in the Center for Environmental Biotechnology, Knoxville, using Illumina HiSeq, 2×250 bp in paired-end mode [43]. Sequencing output summaries are provided in Supp. Table 1.

Sequence reads were quality filtered, trimmed, and normalized as described in the Supplementary information. Processed reads from each sample were assembled separately using IDBA-UD (version 1.1.1) [46] with default settings and the following options: --min_contig 500 -- pre_correction. Reads from site F (Kongsfjorden) were assembled via metaSPAdes (version 3.11) [47] with kmer sizes set to 21, 33, 55, 77, 99, and 127 to find the best assembly. All other samples were assembled using metaSPAdes on the KBase server [48] with the default parameters and following options: minimum contig length of 1000 bp, and kmer sizes of 21, 33, and 55. All samples were also assembled using Megahit [49] on the KBase server using default parameters.

Coverage profiles of assembled unbinned contigs were acquired by mapping trimmed reads (not normalised) to assemblies using BWA [50] and SAMtools [51]. Contigs from each assembly were then binned into metagenome-assembled genomes (MAGs) using MetaBat2 (using each binning strategy) (version 2.12.1) [52], CONCOCT (version 0.4.1) [53] and MaxBin2 (version 2.2.4). MAG collections derived from each binning strategy, from all respective assemblies, were then aggregated using DasTool (version 1.1.0) (Supp. Fig. 1A) [54]. Finally, all MAGs were dereplicated using dRep (version 1.4.3) [55], with the options: an average nucleotide identity (ANI) of 98% was used as cut-off to dereplicate MAGs from the secondary ANI comparison [56], and MAGs >50% complete and <10% contamination were retained. Estimations of completeness and degree of contamination of MAGs were obtained by CheckM (version 1.0.7) [57]. Read mapping to compare relative abundances of read recruitment to MAGs was performed using BBMap [58], with the default settings and ‘minid’ of 0.99 for the minimum identity threshold. Taxonomic affiliations of MAGs were determined with GTDB-Tk [59]. ANI comparisons of MAGs were obtained using JSpeciesWS server based on BLASTN (‘ANIb’) [60] and ANIcalculator [61].

### Gene annotations and in silico analyses of inferred proteins

Calling of genes and annotations were performed via RAST [62]. Functions of predicted proteins of interest were manually checked after searches with BLASTP [63] against the NCBI-nr and SWISS-PROT databases [64] (>30% identity), and the Conserved Domain Database (CDD) [65] (default expect value of 0.01). Functional predictions for proteins were also evaluated using literature searches and the MetaCyc database [66]. Methods for further annotations and protein sequence analyses are described in the Supplementary information.

### MiSeq amplicon sequencing

For amplification of bacterial and archaeal 16S rRNA genes or transcripts (cDNA) from Smeerenburgfjorden sediments, the primers 515F (5’-GTGYCAGCMGCCGCGGTAA-3’) [67] and 806R (5’-GGACTACNVGGGTWTCTAAT-3’) [68] including a 5’-head sequence for 2-step PCR barcoding [69], were used (further details in Supplementary information). Slight variants of these PCR primers 515F and 806R [70] for 16S rRNA genes were used in amplicon sequencing profiling of sediments from Van Keulenfjorden in a previous study, although a standard ‘one-step PCR’ approach was used [40]. Amplicon pools were extracted from the raw sequencing data using the FASTQ workflow in BaseSpace (Illumina) with default parameters. Demultiplexing was performed with the python package demultiplex (Laros JFJ, github.com/jfjlaros/demultiplex) allowing one mismatch for barcodes and two mismatches for linkers and primers. DADA2 [71] was used for demultiplexing amplicon sequencing variants (ASVs) using a previously described standard protocol [72]. FASTQ reads 1 and 2 were trimmed at 220 nt and 150 nt with allowed expected errors of 2. Taxonomy was assigned to 16S rRNA gene/transcript sequences based on SILVA taxonomy (release 138) using the naïve Bayesian classification method as implemented in *mothur* [73]. Amplicon sequence datasets were analyzed with the Rhea pipeline [74] implemented in R (https://www.r-project.org/).

Primers DSR-1762Fmix and DSR-2107Rmix, including a 5’-head sequence for barcoding, were used for amplification of *dsrB*-genes or -transcripts (cDNA) [75] (further details in Supplementary information). Raw reads were then processed as previously described [69, 75], into *dsrB* operational taxonomic units (OTUs) with >99% identity. Classification of DsrB sequences was performed using a combined phylogenetic and naïve Bayesian classification approach as previously described [75].

### Quantitative reverse-transcription PCR

RT-qPCR assays targeting the octaheme cytochrome tetrathionate reductase (*otr*) and *dsrB* genes of MAG AM3-C were performed using the newly-designed primers TetraC-C-F (5’-CACCACGACCTGTCTCGG-3’) and TetraC-C-R (5’-CCCCCTGGAGTTCTTGGT-3’), and Acido-dsrB-F (5’-GGAGAACTATGGGAAGTGGG-3’) and Acido-dsrB-R (5’-GTTGAGGCAGCACGCGTA-3’). Primers 1329-B-F (5’-AACCTTTGGGCGATTTCTCG-3’) and 1329-B-R (5’-GAGAGAGTGGCAACGTGAAC-3’) targeting the DNA-directed RNA polymerase alpha subunit gene of MAG AM3-C were used to examine expression of a housekeeping gene. Details of RT-qPCR assay conditions are presented in the Supplementary information.

### Phylogenetic analyses

A phylogenomic maximum-likelihood tree was created using the IQ-TREE web-server with automatic substitution model selection and ultra-fast bootstrapping (1000×) [76] using an alignment of concatenated protein sequences derived from single copy marker genes retrieved from CheckM [57]. The tree was visualized with iTol [77]. Phylogenetic analysis of 16S rRNA was performed in ARB [78] using the SILVA database release 138 [79], and *dsrAB* sequences were also analysed using ARB using previously described database [36, 75] (Supplementary information). Phylogenetic analyses of all other protein sequences were performed using the IQ-TREE web-server with automatic substitution model selection and ultra-fast bootstrapping (1000×) [76]. For the Complex-Iron-Sulfur-Molybdoenzyme (CISM) tree, query protein sequences were added to a pre-computer alignment of CISM protein sequences [80], using MAFFT using the ‘add full length sequences’ option (--add) [81]. All other proteins sequence alignments were made *de novo* with MUSCLE [82] within Mega6 [83].

### Catalyzed reporter deposition-fluorescence in situ hybridisation (CARD-FISH)

Sediment samples from Svalbard, Portugal and Sweden were fixed with 4% formaldehyde for 3 hrs on ice and stored in PBS:ethanol (1:1) at -20°C using standard procedures [84]. Cells were extracted from sediments using Nycodenz density gradients (Supplementary information). Hybridisations were performed using the 5’-horseradish peroxidase-labeled (HRP) probe Acido-Sva-34-HRP (5’-GACTTATGTCATTGAGGACTCATGCGG-3’) and unlabelled helper probes (5’-GGATAGCCTCGGGAAACCGAGGGTAA-3’) and (5’-TGAGGGGAAAGGCGGGG-3’), or with HoAc1402-HRP (5’-CTTTCGTGATGTGACGGG-3’) with competitor compHoAc1402 (5’-CTTTCGTGACGTGACGGG-3’) [85]. Further details of hybridisation methods and probes are provided in the Supplementary information.

### Sequence and MAG accessions

Metagenomic sequence reads from Van Keulenfjorden and Kongsfjorden samples are available under NCBI-Genbank Bioproject PRJNA493859. Metagenomic sequence reads, and 16S rRNA gene and *dsrB* sequence reads from Smeerenberfjorden samples are available under NCBI-Genbank Bioproject PRJNA623111. Metagenome-assembled genomes are available under NCBI-Genbank Bioproject PRJNA623111, with Biosample accessions SAMN15691661-SAMN15691666.

## Results

### Recovery of novel Acidobacteriota genomes from marine sediments

Metagenomic sequencing and genome binning was performed from DNA extracted and sequenced from sediments originating from three fjords from Svalbard, Norway (Supp. Table 1). Our genome binning strategy based on multiple assemblies and multiple binning algorithms recovered more MAGs with higher completeness, as compared to applying multiple binning approaches based on single assembly approaches (Supp. Fig. 1A and Supp. Fig. 1B). From the dereplicated MAGs (*n*=97), four represented populations of the phylum Acidobacteriota and were chosen for in-depth analyses.

**Figure 1.**
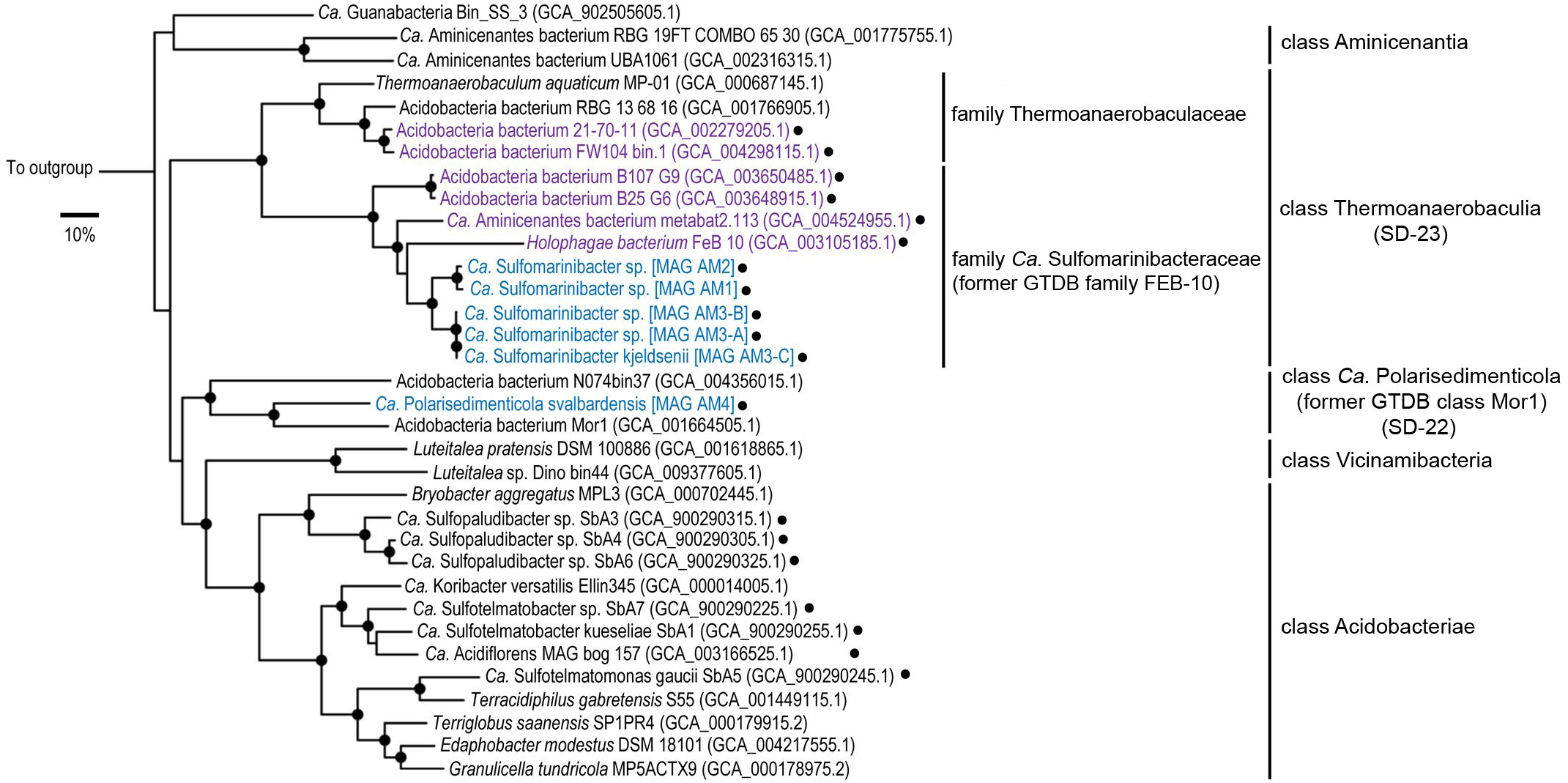
Phylogenomic analysis reveals novel Acidobacteriota taxa in marine sediments. Maximum-likelihood tree of concatenated protein sequences from MAGs and genomes. Single marker genes were retrieved with CheckM. Highlighted in blue are MAGs obtained in this study. Highlighted in purple are *dsrAB*-containing MAGs obtained from the NCBI database from the class Thermoanaerobaculia. The genus *Ca*. Acidiflorens is represented by the most complete MAG (GCA_003166525.1) from the corresponding study [12]. Our phylogenomic analysis showed that one MAG that was previously assigned to *Ca*. Aminicenantes (GCA_004524955.1), recovered from the Bothnian Sea [130], is affiliated with the newly proposed family *Ca*. Sulfomarinibacteraceae. Black dots indicate *dsrAB*-containing genomes/MAGs. Bootstrap values with >90% are indicated with filled black circles on nodes. *Nitrospina gracilis* 3/211 (GCA 000341545.2) was used as an outgroup. The scale bar represents 10% sequence divergence.

Phylogenomic analyses showed three MAGs (AM1, AM2 and AM3-A) affiliated with GTDB family ‘FEB-10’ of the class Thermoanaerobaculia (‘sub-division 23’) (Fig. 1). We included two additional MAGs in our analyses, i.e., AM3-B and AM3-C, that were highly similar to the AM3-A MAG (>98% ANI), but were classified as redundant during MAG dereplication. They encoded enzymes of interest, and were more complete than MAG AM3-A (Table 1). Comparisons of ANI values suggested these MAGs represent three distinct species (<95% ANI) (Supp. Table. 2) [86], all from a novel genus for which we propose the name *Candidatus* Sulfomarinibacter. The MAG AM3-C represents the type species *Ca*. Sulfomarinibacter kjeldsenii (Supp. Table 3). MAG AM4 represents the type species of another novel genus affiliated with the GTDB class ‘Mor1’ (‘sub-division 22’) (Fig. 1), and for which we propose the name *Ca*. Polarisedimenticola svalbardensis (Table 1 and Supp. Table 3).

**Table 1.** Summary of Acidobacteriota MAGs retrieved from Svalbard.

| MAG | MAG (bp) | Estimated<br>genome<br>size (bp) | GC (%) | Complete-<br>ness (%) | Contam-<br>ination<br>(%) | Strain<br>hetero-<br>geneity | Number<br>of<br>contigs | GenBank accession |
| --- | --- | --- | --- | --- | --- | --- | --- | --- |
| *Ca* . Sulfomarinibacter<br>MAG AM1 | 2,426,940 | 3,362,810 | 63.3 | 72.17 | 2.94 | 40 | 666 | JACXVY000000000 |
| *Ca* . Sulfomarinibacter<br>MAG AM2 | 1,835,749 | 3,146,099 | 63 | 58.35 | 7.69 | 43 | 828 | JACXVZ000000000 |
| *Ca* . Sulfomarinibacter<br>MAG AM3-A | 1,779,173 | 2,206,863 | 60.9 | 80.62 | 1.1 | 0 | 127 | JACXWA000000000 |
| *Ca* . Sulfomarinibacter<br>MAG AM3-B | 3,394,205 | 3,824,456 | 60.7 | 88.75 | 4.27 | 20 | 533 | JACXWB000000000 |
| *Ca* . Sulfomarinibacter<br>kjeldsenii MAG AM3-C | 3,921,116 | 4,316,434 | 60.9 | 91.17 | 3.42 | 20 | 783 | JACXWC000000000 |
| *Ca* . Polarisedimenticola<br>svalbardensis MAG AM4 | 3,685,148 | 3,887,504 | 62.2 | 95.73 | 5.98 | 0 | 180 | JACXWD000000000 |

### Marine Acidobacteriota encode the full dissimilatory sulfate reduction pathway

Together, the gene content of the *Ca*. Sulfomarinibacter MAGs suggests they encode a complete canonical dissimilatory sulfate reduction pathway (Fig. 2 and Supp. Table 4). This includes enzymes required for sulfate activation to APS (Sat) and reduction of APS to sulfite (AprAB, QmoABC), and further reduction of sulfite to sulfide (DsrAB, DsrC, DsrMKJOP, DsrN) (Supp. Table 4). Acidobacteriota *dsr* were also found on scaffolds (up to 20 kb) that were not binned into MAGs, yet had highly similar genes and therefore derive from closely related populations, e.g., >99% *dsrB* nucleotide identity (Fig. 3 and Supp. Table 4). The unbinned acidobacteriotal contig ‘ThM_scaffold_807’ harboured all *dsr* on one contig (Fig. 3). The predicted DsrC had two conserved cysteine residues critical for respiratory functioning (Supp. Fig. 2) [87]. Similar to Acidobacteriota MAGs from peatlands and permafrost [2, 12], the marine Acidobacteriota encoded both DsrL and DsrD proteins. DsrL acts as a NAD(P)H:acceptor oxidoreductase for DsrAB [88], while the function of DsrD has not been proven, it is possibly a transcriptional regulator [89]. The DsrL sequences were phylogenetically related to group ‘DsrL-2’ from *Desulfurella amilsiiI*, peatland Acidobacteriota and other subsurface bacteria, and were phylogenetically distinct from group ‘DsrL-1’ of sulfur-oxidizing aerobes (Supp. Fig. 3A) [136]. The DsrL had conserved YRR-motifs in the NAD(P)H substrate-binding domains that are present in the DsrL-2 group, and absent in DsrL-1 of sulfur-oxidizing aerobes (Supp. Fig. 3B) [136].

**Figure 2.**
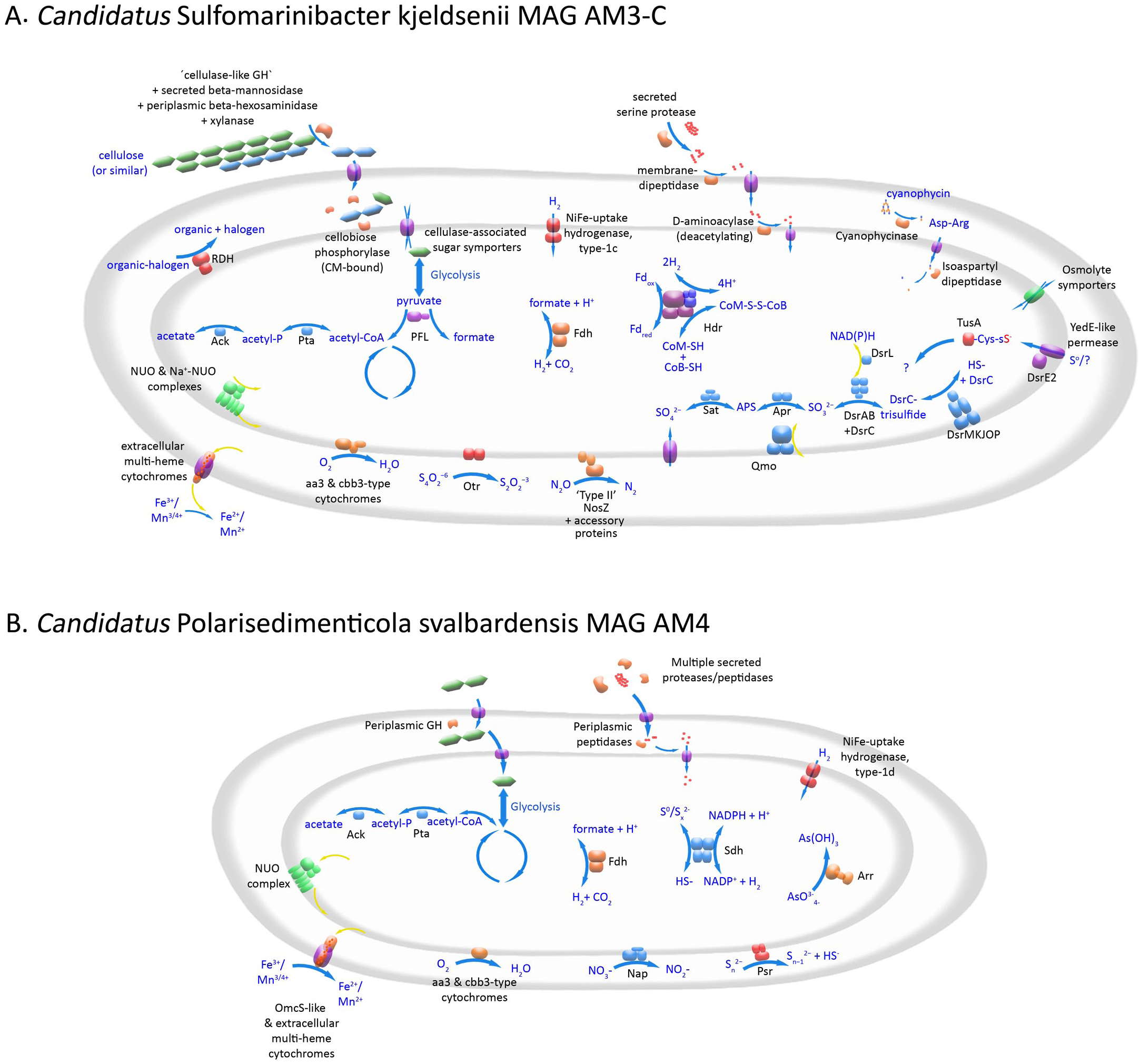
Metabolic model of (A) *Ca*. Sulfomarinibacter kjeldsenii MAG AM3-C and (B) *Ca*. Polarisedimenticola svalbardensis MAG AM4. GH = glycoside hydrolase, RDH = reductive dehalogenase homologous enzyme, Ack = acetate kinase, Pta = phosphotransacetylase, PFL = pyrivate-fomate lyase, FDH = formate dehydrogenase, Hdr = heterodisulfide reductase, NUO = NADH dehydrogenase, Otr = Tetrathionate reductase, NosZ = nitrous oxide reductase, SAT = sulfate adenylyltransferase, APR = adenylylsulfate reductase, Omo = quinone-interacting membrane oxidoreductase complex, Dsr = dissimilatory sulfate reductase, Nap = Periplasmic nitrate reductase, Psr = poylsulfide reductase, Sdh = Sulfhydrogenase complex, TusA = sulfur carrier protein.

**Figure 3.**
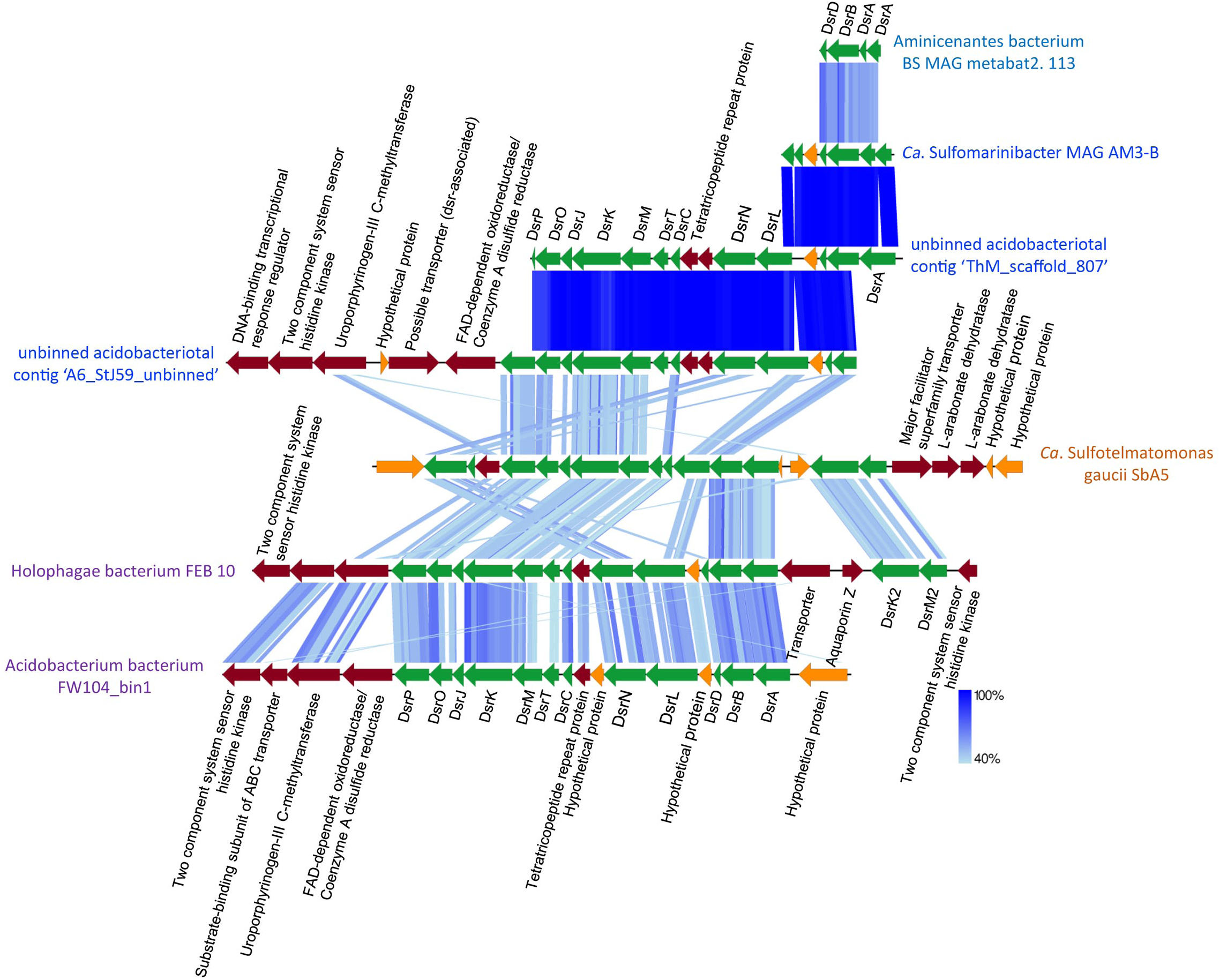
Gene organization of the dsr gene cluster in Acidobacteriota. Scaffold names in blue were retrieved from this study. Scaffold names in purple were derived from best BLASTP hits to sequences from this study. *Ca*. Sulfotelmatomonas gaucii SbAS was retrieved from Hausmann et al. 2018. Green: dsr, dark red: other genes, and orange: hypothetical genes. Shaded blue lines indicate degree of sequence similarity as determined by tBLASTx within EasyFig.

The DsrAB sequences from the novel Acidobacteriota MAGs and unbinned metagenomic contigs are affiliated with the ‘Uncultured family-level DsrAB lineage 9’ within the ‘Environmental supercluster 1’, which is part of the ‘reductive, bacterial-type DsrAB branch’ in the DsrAB tree [36] (Fig. 4). Sequences of ‘lineage 9’ are primarily derived from marine sediments [36]. This lineage is closely related to the ‘Uncultured family-level lineage 8’ that harbours DsrAB sequences from peatland and permafrost derived Acidobacteriota of subdivisions 1 and 3 [2, 12]. We also identified several ‘lineage 9’ *dsrA* and/or *dsrB* sequences in Acidobacteriota MAGs from public databases that derived from marine or groundwater environments (Fig. 4). Herein, we refer to this clade as the ‘Thermoanaerobaculia Dsr lineage’.

**Figure 4.**
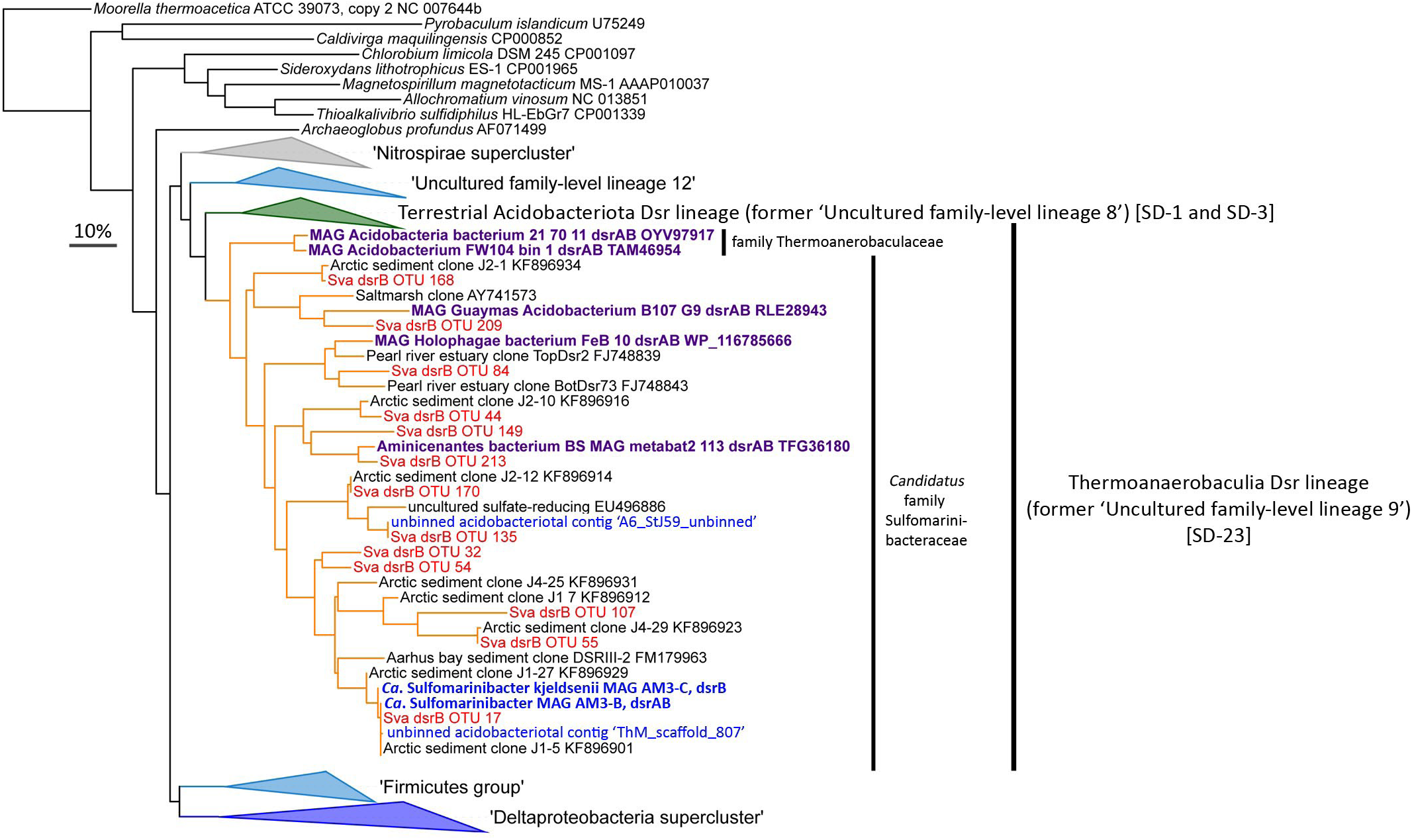
DsrAB uncultured lineage 9 in the DsrAB tree represents members of the Acidobacteriota class Thermoanaerobaculia (sub-division 23). Blue leaves in the DsrAB tree represent MAGs or contigs identified in this study. Red leaves represent the most abundant acidobacteriotal amplicon-derived DsrB sequences identified in this study. Purple leaves represent sequences from MAGs retrieved from public databases. The DsrAB sequences were added to the consensus tree from MOiier et al. 2015 in ARB. SD, pertaining to ’sub-divisions’ of Acidobacteriota. The scale bar represents 10% sequence divergence.

### Marine Acidobacteriota use tetrathionate and potentially also other sulfur cycle intermediates

Several *Ca*. Sulfomarinibacter MAGs encoded c-type cytochromes annotated as octaheme tetrathionate reductases (Otr), which was supported by phylogenetic analysis (Supp. Fig. 4) [90]. The Otr were predicted to be periplasmic and may enable respiration with tetrathionate, a sulfur compound of intermediate oxidation state (‘sulfur cycle intermediate’ (SCI)) [31] (Fig. 2). Transcription of *otr* in Svalbard sediment microcosms with or without tetrathionate additions was analysed by RT-qPCR analysis of mRNA of *otr* of *Ca*. Sulfomarinibacter MAG AM3-C. This showed *otr* was upregulated (1.8-fold) at day 1 although not significantly, and was significantly upregulated (p<0.0488) at day 8 (36-fold). The transcription of *dsrB* appeared lower at both days in tetrathionate-amended microcosms (0.48-0.63-fold), although not significantly (Fig. 5).

**Figure 5.**
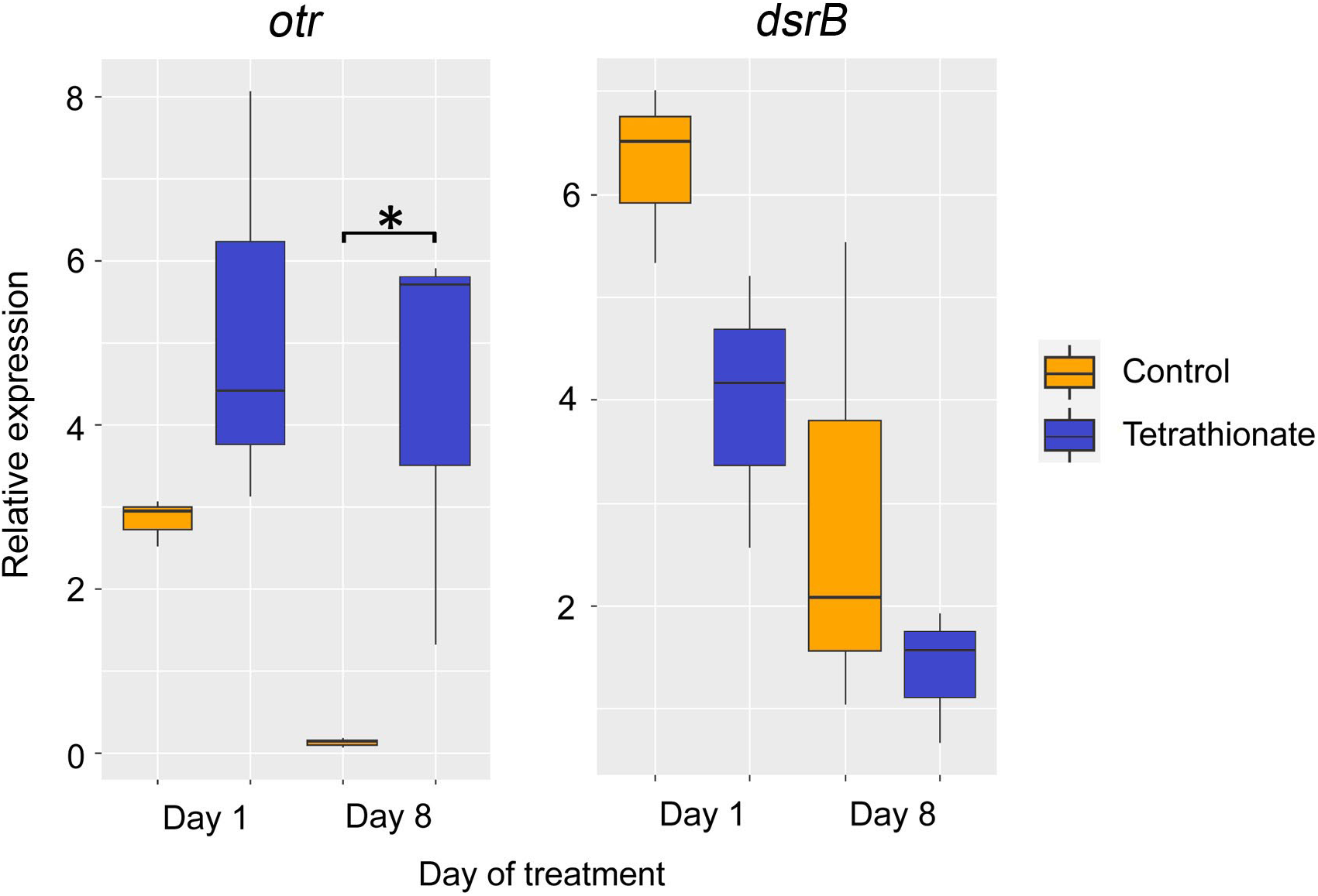
Box plots depicting the expression of *otr* and *dsrB* relative to a house-keeping gene (DNA-directed RNA polymerase, alpha subunit) from *Ca*. Sulfomarinibacter kjeldsenii MAG AM3-C during microcosm experiments with amendments of tetrathionate versus no-amendment controls. Relative expression was determined by RT-qPCR. Expression of otr was significantly higher at day 8 (*p*=0.0488) as determined using a two-tailed T-test, and is indicated by an asterisk. Center lines indicate medians; box limits indicate 25th and 75th percentiles; and whiskers extend 1.5 times the interquartile range from the 25th and 75th percentiles.

‘YTD gene clusters’ encoding sulfur-trafficking rhodonase-like proteins [91] were identified among *Ca*. Sulfomarinibacter MAGs. Genes for YedE-related permease-like proteins, a DsrE2-like protein, a rhodonase-domain containing sulfur carrier TusA, and two conserved hypothetical proteins were present (Supp. Table 4). The TusA sulfurtransferase had conserved Cys□Pro□X□ Pro sulfane sulfur□binding domains (Supp. Fig. 6A). The TusA were phylogenetically most closely related to various TusA from anaerobic Desulfobacterota that are capable of reducing and/or disproportionating inorganic sulfur compounds such as elemental sulfur, sulfite and/or thiosulfate (Supp. Fig. 6B). Together, this suggested *Ca*. Sulfomarinibacter are capable of internal trafficking of sulfur, and may use it to reduce and/or disproportionate inorganic sulfur compounds of intermediate redox states.

The marine Acidobacteriota MAGs encoded several Complex-Iron-Sulfur-Molybdoenzyme (CISM) enzymes that may catalyse redox reactions of sulfur compounds. The *Ca*. Sulfomarinibacter MAG AM3-A encoded a putative tetrathionate reductase (TtrA) (Supp. Fig. 6), and also had an adjacent TtrB (FeS protein) encoded. A *ttrC* encoding a membrane anchor was missing, although the *ttrAB* were situated on the end of the contig and therefore *ttrC* may have been present in DNA that either was not sequenced or was not binned. The Ttr complex may provide an additional means to reduce tetrathionate.

*Ca*. P. svalbardensis MAG AM4 had genes for a CISM subunit A enzyme that phylogenetically affiliated with the polysulfide/thiosulfate reductase clade (‘Psr’) (Supp. Fig. 6). Subunits for PsrABC were encoded in a gene cluster, where the terminal reductase PsrA had a TAT-leader peptide for export from the cytoplasm, PsrB had FeS domains for electron transfer between PsrA and PsrC, and the PsrC subunit was predicted to be membrane-bound. This suggested a periplasm location and that the complex may play a role in respiration of sulfur/polysulfide or thiosulfate. Selenite reductases (SrrA) also phylogenetically affiliate with the polysulfide/thiosulfate reductase clade, but conserved rhodonase-like proteins encoded in the gene neighbourhood of SrrA are thought to be indicative of selenite-reducing organisms [92], but were absent near *psrABC* in MAG AM4.

*Ca*. P. svalbardensis MAG AM4 also harboured a gene cluster encoding four subunits of a sulfhydrogenase complex (Supp. Table 4). Similar to the characterized sulfhydrogenase from *Pyrococcus furiosus*, this included two NiFe hydrogenase subunits, as well as two subunits of anaerobic sulfite reductases [93–95]. These complexes can use elemental sulfur or polysulfides as electron sinks when available [95], or act in reverse as hydrogen-evolving hydrogenases during fermentative growth [96].

### Marine Acidobacteriota may respire additional electron acceptors including metals

All MAGs had gene clusters encoding multi-heme c-type cytochromes with predicted periplasmic or extracellular locations, as well as associated predicted β-barrel proteins (Supp. Table 4). In known metal-reducing and/or -oxidizing bacteria, extracellular and periplasmic cytochromes insert into outer-membrane traversing β-barrel proteins, and transfer electrons through the complexes to/from metals [97, 98]. These gene clusters were syntenous among the MAGs and *Thermoanaerobaculum aquaticum* (Supp. Fig. 7A), a related hot spring-derived isolate that can anaerobically reduce iron- and manganese-oxides [11]. We therefore propose these cytochromes are likely candidates for facilitating the reduction of metal-oxides by *Thermoanaerobaculum aquaticum*, because no other predicted extracellular cytochromes are encoded. We therefore also propose the similar cytochromes in our marine MAGs may also perform this function.

The *Ca*. P. svalbardensis MAG AM4 encoded two additional cytochrome c proteins with similarity to metal-reducing outer-membrane cytochromes (OmcS) from known metal-reducing bacteria, i.e., various Desulfuromonadia (formerly Desulfuromonadales) such as *Geobacter* and *Geopsychrobacter* spp. (Supp. Table 5) [99, 100]. These cytochromes had six heme-binding sites like characterised OmcS, and were also clustered among genes for predicted periplasmic cytochromes and β-barrel proteins (Supp. Fig.7B). They could therefore also potentially exchange electrons with metal oxides (or other insoluble substrates such as humic-like substances, or other cells).

The marine Acidobacteriota MAGs also encoded the potential to reduce oxygen (Fig. 2), nitrous oxide (Supp. Fig. 8), organohalides (Supp. Fig. 9), nitrate, and arsenate (Supp. Fig. 6, Supp. Table 4, and further detailed in Supplementary information).

### Additional energy conserving mechanisms among marine Acidobacteriota

Electron bifurcating heterodisulfide reductase complexes were only encoded in *Ca*. Sulfomarinibacter MAGs (Supp. Table 4). These complexes enable flavin-based redox balancing and formation of low-potential electron carriers (i.e., ferredoxin and/or flavodoxin), and are common among strict anaerobes [101, 102]. A high-molecular-weight cytochrome c3-type protein and a predicted periplasmic location was encoded in *Ca*. Sulfomarinibacter MAG AM1 (Supp. Table 4). These typically act as periplasmic redox hubs to link electron flows between the periplasm and cytoplasm in SRM [103]. All Acidobacteriota MAGs recovered in this study encoded NADH-ubiquinone oxidoreductase (Nuo) complexes required for energy conservation via respiration (Supp. Table 4). Genes for additional sodium-dependent Nuo complexes were also present (Supp. Table 4). Apart from the potential for respiration, some Acidobacteriota MAGs from both *Ca*. Sulfomarinibacter and *Ca*. P. svalbardensis MAG AM4 encoded acetate kinase and phosphate acetyltransferase for fermentation via acetogenesis, or which may act in reverse to facilitate acetate consumption (Supp. Table 4).

### Marine Acidobacteriota use diverse nutrient and electron sources

The *Ca*. Sulfomarinibacter AM3 MAGs encoded predicted cellulase A enzymes with signal peptides for export from the cytoplasm (Fig. 2). They were phylogenetically affiliated with cellulase A from various anaerobic degraders of cellulose and/or plant-derived polysaccharides (Supp. Fig. 10). A cellobiose phosphorylase was encoded in *Ca*. S. kjeldsenii MAG AM3-C, and had relatively high amino acid identity (63%) to a characterized cellobiose phosphorylase from *Thermotoga neapolitana* [104]. These enzymes catalyse phosphorolysis of cellobiose to а-D-glucose 1-phosphate (G1P) and D-glucose, thereby saving an ATP before entering glycolysis, and are typically used by anaerobic cellulose-degraders [105]. This suggests these organisms have the capacity to anaerobically degrade cellulose, a derivative of cellulose, or a structurally similar compound. Overall, the marine *Ca*. Sulfomarinibacter MAGs encoded few genes for glycoside hydrolases or other carbohydrate active enzymes, i.e., 0.47-0.75% of protein encoding genes encoded glycoside hydrolases (further detailed in Supplementary information) (Supp. Table 6). The *Ca*. P. svalbardensis MAG AM4 also encoded few glycoside hydrolases (0.54% of protein encoding genes), with none predicted to be exported to the extracellular environment, and a single endo-1,4-beta-xylanase predicted to be periplasmic (Supp. Table 4).

Genes for cyanophycinases among *Ca*. Sulfomarinibacter MAGs indicated they may utilize the storage compound cyanophycin as a nutrient (Supp. Table 4). The cyanophycinases had Secretion-signal peptides (Sec-) for export from the cytoplasm, indicating they act on an external substrate and not an internally stored compound. Accordingly, no genes for cyanophycin synthetases were found. An isoaspartyl dipeptidase was encoded in *Ca*. S. kjeldsenii MAG AM3-C, which may enable utilization of the products released by the cyanophycinase, i.e., a dipeptide of aspartate and arginine (Supp. Table 4). The capacity to catabolically degrade aspartate and arginine was also encoded (Supp. Table 4).

The *Ca*. Sulfomarinibacter MAG AM3-C may degrade extracellular proteins using two predicted secreted proteases, as well as adjacently encoded peptidases predicted to be membrane-bound (Supp. Table 4). The *Ca*. P. svalbardensis MAG AM4 harboured numerous genes for proteases/peptidases (*n*=7) that were predicted to be secreted, strongly indicating these bacteria use proteins as nutrients (Supp. Table 4).

Membrane-bound NiFe uptake-hydrogenases were encoded by both *Ca*. Sulfomarinibacter and *Ca*. P. svalbardensis MAGs (Supp. Table 4). These may be important for oxidizing environmental hydrogen. The *Ca*. Sulfomarinibacter MAGs encoded ‘type-1c’ NiFe hydrogenases typically found in obligate anaerobes and that are thought to be oxygen sensitive (Supp. Fig. 11) [106]. The *Ca*. P. svalbardensis MAG AM4 encoded a ‘type-1d’ NiFe hydrogenase, which are typically found in aerobes and facultative anaerobes (Supp. Fig. 11) [106]. Inspection of best BLASTP hits from the NCBI-nr database to the *Ca*. Sulfomarinibacter NiFe hydrogenase sequences identified various sequences previously shown to be expressed in tidal flat sediments [107]. Formate dehydrogenases encoded among MAGs of both *Ca*. Sulfomarinibacter and *Ca*. P. svalbardensis also suggested formate may be used as an electron donor (Supp. Table 4).

### Adaptations to marine environments

Comparative genomics with seven *dsr*-harbouring Acidobacteriota MAGs from peatland soil [2] suggested the marine Acidobacteriota encoded unique adaptations to marine settings (Supplementary information) (Supp. Fig. 12). These included various predicted transporters/symporters and pumps for ions (e.g., sodium and potassium) and metals/metalloids (e.g., zinc and arsenic) that were unique to the marine MAGs. Genes for a sodium-translocating NADH-quinone oxidoreductase complex, which are used by various marine microorganisms to support respiration and cellular homeostasis [108], were only present in marine MAGs. Symporters for the osmolytes proline, glutamate and glycine, were also only present in marine MAGs.

### Acidobacteriota are abundant, active and diverse in marine sediments

Amplicon sequencing of 16S rRNA genes revealed Acidobacteriota had an average relative abundance of 4.5±2.2% in Smeerenbergfjorden sediments (Supp. Fig. 13 and 14), which have high sulfate reduction rates (reaching around 100 nmol SO_4_ ^-2^ cm^-3^ d^-1^ around 5 cmbsf) [38, 41, 109]. Thermoanaerobaculia-affiliated sequences were the most dominant of any Acidobacteriota, and reached the most abundant (11%) genus-level clade of *Bacteria* at 31 cmbsf in Station J (2016). The same clade was on average the fourth most abundant genus-level clade in the same core (averaged 4.5±2.8%). 16S rRNA transcripts of Acidobacteriota were below 0.5% relative abundances in the surface sediments (0–1 cmbsf) of Smeerenbergfjorden cores (Supp. Fig. 14). At station GK, Acidobacteriota 16S rRNA transcripts reached 6% relative abundance at 15 cmbsf (Supp. Fig. 14). We also examined Acidobacteriota 16S rRNA genes from metal-rich Van Keulenfjorden sediments from a previously published study [40]. This showed *Ca*. Polarisedimenticola related sequences were the most prominent Acidobacteriota, reaching 1.5%, and averaging 1.1±0.21% of communities in four cores (Supp. Fig. 14). Members of the Thermoanaerobaculia were in much lower abundances (0.3±0.2% average overall), although they reached 1.1% in deeper sections of core AB. Mapping of metagenomic reads to the Acidobacteriota MAGs supported the general distribution trends from 16S rRNA amplicon analyses, i.e., that Thermoanaerobaculia were abundant in Smeerenbergfjorden sediments and *Ca*. Polarisedimenticola were more abundant in Van Keulenfjorden sediments (Supp. Table 7, and further detailed in Supplementary information).

Phylogenetic analysis of 16S rRNA genes from Smeerenbergfjorden sediment (Supp. Fig. 15) and examination of Acidobacteriota 16S rRNA sequences in the SILVA database (Supp. Fig. 16) revealed diverse Acidobacteriota sequences from marine sediments. It also revealed that Thermoanaerobaculia (sub-division 23) and *Ca*. Polarisedimenticolia (sub-division 22) sequences are the most prominent Acidobacteriota lineages in marine sediments in general (further detailed in Supplementary information).

Sequencing of *dsrB* genes and transcripts from Smeerenburgfjord sediments revealed the Acidobacteriota *dsrB* averaged 13±6.6% of all *dsrB* (DNA-derived) sequences, and 4±2 % of *dsrB-* transcripts (cDNA-derived) (Supp. Fig. 17). Acidobacteriota *dsrB* sequences were the second most abundant group after Desulfobacterota *dsrB*, which dominated the sediments and averaged 75±6% in relative abundance (Supp. Fig. 17). Acidobacteriota *dsrB* reached a maximum of 19% at station GK and 31% at station J. The most abundant Acidobacteriota *dsrB-*OTU-17 was 100% identical (over 321 nucleotides) to *dsrB* from *Ca*. Sulfomarinibacter AM3-B MAG (Fig. 4). Amplicon-derived DsrB sequences that affiliated with the DsrB from marine Acidobacteriota MAGs were phylogenetically diverse and spread through-out the ‘Thermoanaerobaculia Dsr clade’ (Fig. 4).

### Description of novel Acidobacteriota Candidatus taxa

Based on their unique phylogeny, predicted metabolic properties, CARD-FISH visualized cells of Thermoanaerobaculia (thin rods present in three different sites, see Supp. Fig. 18) and relatively complete MAGs, we propose the following new *Candidatus* taxa of Acidobacteriota (Supp. Table 3):

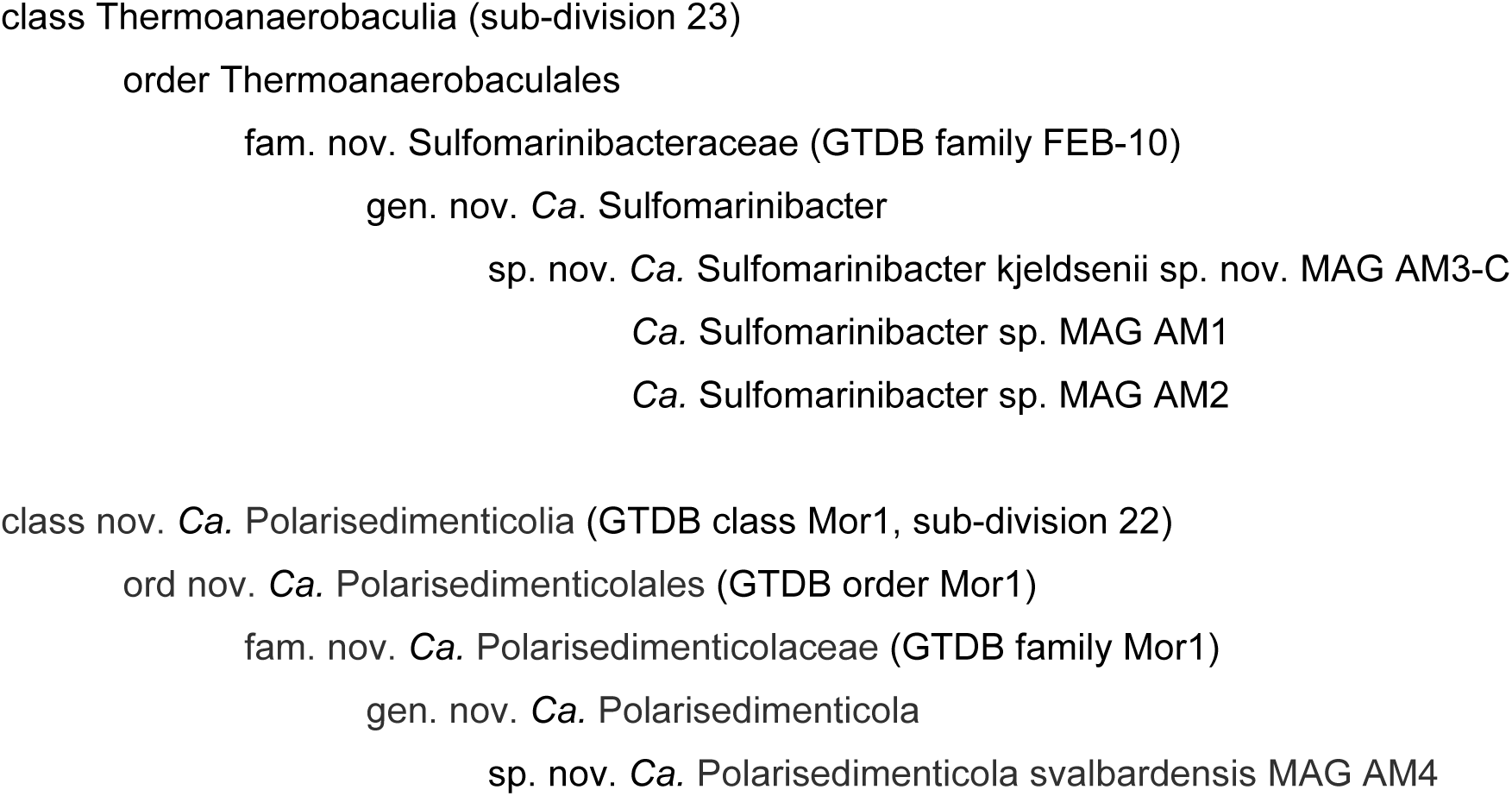

## Discussion

This study provides the first insights into the genomes and metabolic potential of abundant Thermoanaerobaculia from marine sediments, and new insights into the metabolisms of *Ca*. Polarisedimenticolia (sub-division 22, or Mor1). Most notably, we revealed that MAGs from both of the major lineages of Acidobacteriota from marine sediments have capabilities to dissimilate various inorganic sulfur compounds.

Genes for the full dissimilatory sulfate reduction pathway provided the first direct link between genomes of marine sediment Acidobacteriota and DsrAB sequences of the previously undescribed ‘Uncultured family-level lineage 9’ clade (here named ‘Thermoanaerobaculia Dsr lineage’). In addition to being abundant and actively transcribed in Svalbard sediments as shown here, *dsrB* sequences of this lineage often constitute a prominent fraction of *dsrB*-harbouring communities in various sediments, e.g., making around 8–14% of *dsrB* sequences from Aarhus Bay sediments [110, 111], around 5% of *dsrB* sequences in Baltic Sea sediments [112], and up to 15-25% of sequences in sediments from various cores from the Greenland coast [37]. Together, this indicates Acidobacteriota are a widespread and prominent group of inorganic sulfur-dissimilating microorganisms in marine sediments.

While enzymes of the dissimilatory sulfate reduction pathway are widely used for anaerobic reduction of sulfite/sulfate [35], some organisms can use them in reverse for the oxidation of reduced sulfur compounds [113], or for disproportionation of sulfur compounds [114, 115]. Because no enzymes are currently known that distinguish these different metabolisms, discerning sulfur metabolisms based on genomic data requires careful interpretation [114, 115]. For instance, the *Ca*. Sulfomarinibacter MAGs encoded DsrL, which was previously thought to be exclusively found in sulfur-oxidizing bacteria [88]. However, recent work showed DsrL can function in a reductive manner in biochemical assays [88] [136], and was highly expressed during reductive sulfur- and thiosulfate-respiration by *Desulfurella amilsii* [88, 116]. The DsrL of *Ca*. Sulfomarinibacter contained putative NADP(H)-binding domain structures that may enable coupling of NADPH as electron donor to sulfite reduction [136], as well as phylogenetic relatedness with DsrL of *Desulfurella amilsii*. Together, this indicates the DsrL of *Ca*. Sulfomarinibacter has potential to facilitate a reductive pathway.

The *Ca*. Sulfomarinibacter MAGs encoded rhodonase-like TusA and DsrE2, which act as sulfur-trafficking proteins in reverse-Dsr harbouring sulfur-oxidizing bacteria, i.e., they help deliver sulfur to DsrABC for oxidation [117]. Interestingly, the ‘YTD gene clusters’ that encode these enzymes are also common in genomes of anaerobic elemental sulfur-reducing and/or -disproportionating bacteria that have Dsr, and are suggested to be genetic indicators for disproportionation potential among these anaerobes [91]. The TusA proteins from *Ca*. Sulfomarinibacter were most closely related to TusA from various anaerobic sulfur-reducing and -disproportionating Desulfobacteriota (Supp. Fig. 6). This suggested *Ca*. Sulfomarinibacter could reduce and/or disproportionate elemental sulfur, or possibly other sulfur compounds that can be trafficked by TusA, like thiosulfate [118]. Indeed, the ability to disproportionate sulfur compounds is common among sulfate-reducing Desulfobacteriota [119]. Elemental sulfur is often the most abundant sulfur cycle intermediate (SCI) in marine sediments [120], and was measured in sediments from Smeerenbergfjorden up to 0.15 wt % of total sulfur [39]. Overall, the gene content of *Ca*. Sulfomarinibacter MAGs indicated flexible dissimilatory sulfur metabolisms that may be dictated by and/or switch under different biogeochemical and redox conditions.

Results indicated *Ca*. Sulfomarinibacter likely use the dissimilatory sulfate reduction pathway in a reductive direction in most depths of the sediments studied. Firstly, Acidobacteriota were relatively abundant and expressed *dsrB* in deeper (>15-75 cmbsf), strictly anoxic sediment layers of Smeerenbergfjorden. These sediments lack electron acceptors that could sustain these abundant populations growing via biological oxidation of sulfides, i.e., oxygen, nitrate or oxidized metals [31, 121]. In Station J sediments, oxygen and nitrate are depleted within millimetres-to-centimetres of the surface [122, 123], and sulfide oxidation facilitated by Fe(III) is negligible [41]. An alternative possibility is that cryptic biogeochemical cycling could sustain sulfide oxidation, i.e., fast consumption and production of low concentrations of sulfides and oxidants [124]. Nevertheless, it remains unproven whether biological sulfide oxidation occurs in deep sediments that lack measurable concentrations of required oxidants [28]. On the other hand, the relative abundances of Acidobacteriota peaked in subsurface zones around 5 cmbsf in Station J sediments, where sulfate reduction rates also peak [41, 109]. In another study, Acidobacteriota 16S rRNA gene relative abundances were also highly correlated with sulfate reduction rates in sediments from Greenland [37]. These associations therefore point toward an active role in the reduction and/or disproportionation of sulfur compounds of various oxidation states by *Ca*. Sulfomarinibacter in marine sediments.

Our results also suggested marine Acidobacteriota have potential to reduce various inorganic sulfur compounds independent of the Dsr pathway. Our tetrathionate-amended microcosm experiment suggested that *Ca*. Sulfomarinibacter use tetrathionate as an electron sink via cytochromes, which supports the roles of these enzymes in tetrathionate reduction within *in situ*-like conditions. This is noteworthy because these enzymes were only previously shown to perform this function during biochemical assays [125], i.e., their utilization under *in situ*-like conditions was unknown. The ability to utilize SCI, e.g., tetrathionate or elemental sulfur/polysulfides/thiosulfate, could be important in sediment zones where SCI might be generated from sulfides reacting with available oxidants [41].

The *Ca*. Sulfomarinibacter MAGs indicated they could respire oxygen using terminal cbb3- or aa3-type cytochromes, although we speculate these may instead be used for defence against oxygen because they encoded many characteristics of obligate anaerobes (Supplementary discussion). We also hypothesize that the different redox metabolisms of the two predominant Acidobacteriota groups in Svalbard sediments, i.e., the *Ca*. Sulfomarinibacter and *Ca*. Polarisedimenticola, may explain their different abundances among fjords with different biogeochemical properties (Supplementary discussion). That is, the *Ca*. Sulfomarinibacter may be adapted to low redox environments, and are thus more abundant in the reduced (visibly black), sulfidic subsurface sediments of Smeerenburgfjorden. In comparison, the *Ca*. P. svalbardensis MAG had additional genes to utilize high-potential electron acceptors such as oxygen, nitrate and oxidized metals, and may be better adapted to the more high redox, metal-rich sediments of Van Keulenfjorden (visibly reddish-orange).

If members of the *Ca*. Sulfomarinibacter are indeed SRM, a question arises regarding how they co-exist with dominant sulfate-reducing Desulfobacterota populations, as both apparently use hydrogen, acetate or formate as substrates. However, we identified genes for use of several organic substrates that may enable *Ca*. Sulfomarinibacter to occupy a distinct nutrient niche. Complex carbohydrates such as cellulose (or structurally similar compounds) could be used. Carbohydrates are not used by most known isolated Desulfobacterota SRM [35]. Plant-derived molecules could stem from terrestrial run-off, which is a major source of organic carbon to arctic sediments [126, 127] and to coastal marine systems in general [128]. Additionally, various marine algae are known to produce cellulose [129]. The predicted ability to utilize cyanophycin could also facilitate a unique nutrient niche. Cyanophycin is a multi-L-arginyl-poly-L-aspartic acid, commonly produced by cyanobacteria as a storage compound [130, 131]. Indeed, few organisms are known to use cyanophycin anaerobically [132], and no anaerobes are known from marine sediments.

*Ca*. Polarisedimenticola svalbardensis appeared to have a high propensity for the degradation of proteins, which was indicated by a suite of predicted secreted peptidases. A related Mor1 Acidobacteriota genome (GCA_001664505.1) (Fig. 1) was recovered as a bacterial co-inhabitant of a cyanobacterial enrichment culture from seawater, suggesting it used organic material/necromass from the primary-producing cyanobacterium [133]. *Ca*. Polarisedimenticola may therefore contribute to protein degradation in marine sediments, where proteinaceous organics comprise a large proportion (∼ 10%) of available organic matter [134].

In summary, the genome-encoded dissimilatory sulfur metabolisms and the high abundances and activity of *Ca*. Sulfomarinibacter in the sulfidic zones of Svalbard sediments, suggested these novel Acidobacteriota of the class Thermoanaerobaculia (sub-division 23) are important players in the biogeochemical sulfur cycles of the sediments. Our data also indicated that *Ca*. Sulfomarinibacter thrive largely via anaerobic metabolisms with the capability to use various other electron acceptors with different redox potentials, including biogeochemically relevant metal-oxides. Additionally, we show that *Ca*. Polarisedimenticola svalbardensis, a member of a different class of Acidobacteriota (sub-division 22), has the genetic potential for protein degradation and for metabolisms driven by high redox potential electron acceptors such as oxygen, nitrate and metal-oxides.

## Acknowledgements

This research was supported by the Austrian Science Fund (FWF grants P29426 to KW and P25111-B22 to AL) and the MetaBac Research Platform of the University of Vienna. We thank Captain Stig Henningsen of MS Farm during Svalbard expeditions. We particularly thank Bo Barker Jørgensen, Alexander Michaud and Susann Henkel for organising the 2016 and 2017 Svalbard expeditions, and all members of Svalbard expeditions for help with sample collection, especially Claus Pelikan. We thank the Biomedical Sequencing Facility (BSF) Vienna for sequencing of metagenome samples, the Joint Microbiome Facility (JMF) of the Medical University of Vienna and the University of Vienna, and Microsynth for sequencing amplicons. We specifically thank Jasmin Schwarz, Gudrun Kohl and Petra Pjevac from the JMF for assisting with amplicon sequencing. We thank Marc Mussman and Stefan Dyksma for providing the hydrogenase database. We are grateful to Bernhard Schink for help with Latin naming of taxa.

## Contributions

KW and AL conceived the study. KW, JB, KGL and AL collected samples. MF and KW performed microcosm experiments. MF performed DNA/RNA extractions and PCRs for amplicon sequencing. KW, MF and JB performed bioinformatic analyses of metagenomic and amplicon data, with support from CH, BH and TR. MF and BH analysed amplicon sequencing data. MF designed and performed RT-qPCR experiments. KW and MF interpreted genomic data. MF and KW performed CARD-FISH. MF, KW and AL wrote the manuscript, with contributions from all authors.

## Conflict of interest

The authors declare no conflicts of interest.

## Figure captions

**Supplementary Figure 1. A)** Outline of the metagenomic binning strategy. **B)** Plot of completeness of MAGs (CheckM). Comparisons are derived from binning from single assemblies (IDBA or MetaSpades or Megahit), versus binning from multiple assemblies of each sample (outlined in panel A).

**Supplementary Figure 2**. Alignment of dissimilatory DsrC cysteine motifs. Sub-section (C-terminus) of alignment of DsrC proteins, showing two conserved cysteine residues (dark purple) that are present in dissimilatory versions of the enzymes.

**Supplementary Figure 3. A)** Phylogenetic tree of DsrL proteins. The sequences from MAGs recovered in this study are highlighted in blue. Other Acidobacteriota DsrL are highlighted in purple. Bootstrap values >50% are presented on nodes as black-filled circles. The scale bar represents 20% sequence divergence. **B)** Alignment of DsrL proteins. A subsection of the whole DsrL alignnment is shown to highlight YRR amino acids for putative NAD(P)H-binding domain.

**Supplementary Figure 4**. Phylogenetic tree of multiheme cytochrome protein sequences. Sequences from MAGs recovered in this study are highlighted in blue. Sequences from other Acidobacteriota are highlighted in purple. Reference sequences were retrieved from Kern et al., 2011, and from best BLASTP hits to our MAG-derived sequences. Functional assignments are labelled at the end of each leaf label. NrfA = respiratory cytochrome c nitrite reductase, Onr = octaheme cytochrome c nitrite reductase, Hao/Hzo = octahaem hydroxylamine oxidoreductase/hydrazine oxidoreductase, MccA = cytochrome c sulfite reductase, and Otr = octaheme tetrathionate reductase. Genbank accessions are presented in parentheses. The scale bar represents 50% sequence divergence.

**Supplementary Figure 5. A)** Phylogenetic tree of TusA proteins. The sequences from MAGs recovered in this study are highlighted in blue. The orange branch indicates TusA proteins from anaerobic organisms known to reduce or disproportionate sulfur cycle intermediates and that had TusA related to the Aciobacteriota TusA. Descriptions of sulfur metabolisms related to reduction or disproportionation of sulfur cycle intermediates are presented in parenthesis for TusA related to TusA from MAGs recovered in this study. Bootstrap values >50% are presented on nodes as black-filled circles. The scale bar represents 20% sequence divergence. **B)** Alignment of TusA proteins from marine Acidobacteriota showing Cys□Pro□X□Pro sulfane sulfur□binding domains.

**Supplementary Figure 6**. Phylogenetic tree of complex iron–sulfur molybdoenzyme (CISM) family proteins. The sequences from the MAGs recovered in this study are highlighted in blue. Reference sequences were obtained from Duval et al., 2008, as well as selected additional sequences. Bootstrap values >90% are presented on nodes as black-filled circles. The scale bar represents 50% sequence divergence.

**Supplementary Figure 7. A)** Schematic of gene organisation and synteny of extracellular cytochrome-rich genomic loci among Acidobacteriota MAGs (AM1, AM3-C and AM4) and *Thermoanaerobaculum aquaticum* (T. aq.). **B)** Schematic of gene organisation of genomic loci encoding OmcS-like proteins in MAG AM4. Shaded blue lines indicate degree of sequence similarity as determined by tblastx within EasyFig (Sullivan et al., 2011). Subcellular location predictions and number of heme-binding sites (CXXCH) are indicated in parentheses. SEC-peptides for Sec secretion systems were searched in proteins with ‘unknown’ location predictions using PRED-TAT (Bagos *et al*., 2011).

**Supplementary Figure 8**. Phylogenetic tree of nitrous oxide reductases (NosZ). Sequences from MAGs recovered in this study are highlighted in blue. The NosZ from MAG AM1 was omitted due to short sequence length, although it was most similar to the NosZ from AM3-B and AM3-C (>90% amino acid identity from 190 amino acids). Sequences from other Acidobacteriota are highlighted in purple. Clade of ‘type I NosZ’ = blue, and clade of ‘type II NosZ’ = red. Reference sequences were retrieved from the top 50 best BLASTP hits to the NosZ from MAG AM3-C were included. Genbank accessions are presented in parentheses. Black circles on nodes represent bootstraps values >90%. The scale bar represents 20% sequence divergence.

**Supplementary Figure 9**. Phylogenetic tree of reductive dehalogenase homolog A (RdhA) proteins. The sequence from the MAG recovered in this study are highlighted in blue. Reference sequences were obtained from Hug et al., 2013, and the top 10 best BLASTP hits to the RdhA from MAG AM3-C were also included. The RdhA sequence of MAG AM1 was not included due to the truncated protein sequence, although it was most similar to the RdhA from MAG AM3-C (>87% amino acid identity from 120 amino acids). The scale bar represents 50% sequence divergence.

**Supplementary Figure 10**. Phylogenetic tree of cellulase A-like proteins. Sequences from MAGs recovered in this study are highlighted in blue. Sequences from other Acidobacteriota are highlighted in purple. Sequences from genera or species known to perform cellulose degradation are highlighted in green. Reference sequences were retrieved from the top 50 best BLASTP hits to the cellulase A from MAG AM3-C. Genbank accessions are presented in parentheses. The tree was rooted with the cellulase A of *Bacillus subtilis*. Black circles on nodes represent bootstraps values >90%. The scale bar represents 20% sequence divergence.

**Supplementary Figure 11**. Phylogenetic tree of [NiFe]-hydrogenase large subunit proteins. The sequences from the MAGs recovered in this study are highlighted in dark blue. Sequences from other Acidobacteriota are highlighted in purple. Sequences from PCR-derived amplicons from tidal flat sediments (Dyksma et al., 2018) are highlighted in light blue. Reference sequences were derived from best BLASTP hits from NCBI-nr database. Hydrogenase ‘types’ were determined using HydDB (Søndergaard et al., 2016). Black circles on nodes represent bootstraps values >90%. The scale bar represents 20% sequence divergence.

**Supplementary Figure 12**. Comparisons of COG classifications of proteins representing unique ortholog groups (OGs) from marine versus terrestrial dsr-harbouring Acidobacteriota. OGs unique to each group of genomes were determined using OrthoFinder (Emms and Kelly 2019). Proteins were compared from the six MAGs recovered in this study, versus proteins from the seven MAGs recovered by Hausmann et al., 2018. Letters in parenthesis represent standard COG codes.

**Supplementary Figure 13**. Microbial community composition of Smeerenburgfjorden sediments. Relative abundance of 16S rRNA ASVs is derived from amplicon sequencing of the 16S rRNA gene. Taxa that are less abundant than 1% or are unclassified are shown in grey. Depth is shown in centimeters below seafloor (cmbsf) for A) Station J, B) Station GK, and C) Station GN.

**Supplementary Figure 14**. Relative abundances of 16S rRNA and DsrB genes and transcripts for Svalbard sediments. A) Acidobacteriota, B) Thermoanaerobaculia, C) *Ca*. Polarisedimenticola (SD-22), *Ca*. Sulfomarinibacter ASV-2257, E) Thermoanaerobaculia DsrB (uncultured family-level lineage 9). Smeerenburgfjorden stations GK, J and GN were sampled in June 2017, and J16 was sampled in July 2016. Replicate cores from Van Keulenfjorden stations AB and AC are derived from Buongiorno et al., 2019.

**Supplementary Figure 15**. Phylogenetic tree of 16S rRNA gene sequences. Red leaves are from all acidobacteriotal amplicon-derived sequences (ASVs) retrieved in this study from Smeerenbergfjorden, Svalbard. The red dot denotes the most abundant ASV in the dataset. The orange leaf represents the 16S rRNA sequence recovered from an acidobacteriotal metagenome-assembled genome (this study). Blue leaves represent sequences derived from marine environments and present in the SILVA database v138. Green leaves represent cultivated *Acidobacteriota*. SD = ‘sub-division’, and are numbered as per SILVA database (v138). The tree was built as a consensus of three maximum-likelihood methods (see Materials and Methods). The scale bar represents 10% sequence divergence.

**Supplementary Figure 16**. Sankey diagram of taxonomic breakdown of marine sediment derived Acidobacteriota 16S rRNA genes from the SILVA database (v138 NR).

**Supplementary Figure 17**. Community compositions of *dsrB*-harbouring microorganisms in Svalbard sediments. **A)** Compositions determined from *dsrB*-gene (DNA) amplicon sequencing. **B)** Compositions determined by *dsrB*-transcript (cDNA) amplicon sequencing. Groups with relative abundances <1% are grouped as ‘other’.

**Supplementary Figure 18**. CARD-FISH images of Acidobacteriota from marine sediments. Sediment locations and probes used are listed above panels A-E. Panels with blue cells are DAPI stained, panels with green cells are CARD-FISH hybridised cells from corresponding fields of view. White scale bars represent 10 µm.

## Notes

### Competing Interest Statement

The authors have declared no competing interest.

