## Supplementary Tables 1-6 for "Novel taxa of Acidobacteriota involved in seafloor sulfur cycling"

**Supplementary Table 1**. Metagenomic sample summary and information.

| Metagenomic sample origin* | General biogeochemical type | Sample site coordinates | Illumina read pairs | Read coverage normalised | Lab. |
|---|---|---|---|---|---|
| Smeerenburgfjorden; Station J, 0-4 cmbsf | Ferruginous/sulfidic | 79° 42.83N, 11° 05.10E | 45,996,553 | | Vienna |
| Smeerenburgfjorden; Station J, 10-15 cmbsf | Sulfidic | 79° 42.83N, 11° 05.10E | 73,362,243 | | Vienna |
| Smeerenburgfjorden; Station J, 17-20 cmbsf | Sulfidic | 79° 42.83N, 11° 05.10E | 49,176,577 | | Vienna |
| Smeerenburgfjorden; Station J, 59 cmbsf | Sulfidic | 79° 42.83N, 11° 05.10E | 56,792,638 | | Vienna |
| Smeerenburgfjorden; Station J, 5-10 cmbsf, microcosm with tetrathionate + molybdate | Sulfidic | 79° 42.83N, 11° 05.10E | 99,662,741 | | Vienna |
| Smeerenburgfjorden; Station J, 5-10 cmbsf, microcosm with thiosulfate + molybdate | Sulfidic | 79° 42.83N, 11° 05.10E | 53,175,346 | | Vienna |
| Smeerenburgfjorden; Station J, 5-10 cmbsf, microcosm with tetrathionate | Sulfidic | 79° 42.83N, 11° 05.10E | 197,828,199 | to 100x | Vienna |
| Van Kuelenfjorden; Station AC, 18 cmbsf | Ferruginous/ manganous | 77°32.260’ N, 15°39.434’ E | 167,411,750 | to 100x | Vienna |
| Van Kuelenfjorden; Station AB, 0-5 cmbsf | Ferruginous/ manganous | 77°35.249’ N, 15°05.121’E | 98,211,882 | to 100x | Knoxville |
| Kongsfjorden; Station F, 0-5cmbsf | Ferruginous/ manganous | 78°55.075’ N, 12°15.929’ E | 75,827,490 | to 100x | Knoxville |

* all samples taken in July 2016.

**Supplementary Table 2**. Average nucleotide identities (ANI) and percentage aligned [%] of MAGs.

A) Calculated using BLAST using JSpeciesWS.

| | AM1 | AM2 | AM3-A | AM3-B | AM3-C | AM4 |
|---|---|---|---|---|---|---|
| **AM1** | | 94.36 [39.02] | 74.28 [37.35] | 74.82 [51.43] | 74.54 [58.93] | 64.64 [11.90] |
| **AM2** | 94.09 [54.74] | | 75.68 [31.00] | 77.09 [47.22] | 76.86 [54.47] | 65.16 [12.16] |
| **AM3-A** | 73.93 [45.69] | 75.29 [27.42] | | 98.91 [76.99] | 98.84 [84.02] | 64.62 [10.30] |
| **AM3-B** | 74.77 [35.00] | 77.17 [23.07] | 97.72 [42.80] | | 98.41 [71.47] | 64.62 [8.81] |
| **AM3-C** | 74.48 [35.25] | 76.89 [23.71] | 97.42 [39.90] | 98.07 [62.28] | | 64.58 [8.79] |
| **AM4** | 64.73 [7.57] | 65.06 [5.20] | 64.22 [5.88] | 64.83 [8.14] | 64.52 [8.85] | |

B) Calculated using ANIcalculator (https://ani.jgi.doe.gov/html/home.php?).

| | AM1 | AM2 | AM3-A | AM3-B | AM3-C | AM4 |
|---|---|---|---|---|---|---|
| **AM1** | | 96.01 [44] | 74.86 [24] | 76.43 [25] | 75.8 [25] | 68.46 [03] |
| **AM2** | 96.06 [34] | | 76.96 [15] | 79.29 [17] | 78.63 [18] | 69.07 [02] |
| **AM3-A** | 74.79 [17] | 77.4 [13] | | 99.22 [28] | 98.48 [25] | n.a.* |
| **AM3-B** | 76.58 [34] | 79.49 [31] | 97.43 [56] | | 99.13 [47] | 68.69 [03] |
| **AM3-C** | 75.81 [39] | 79.04 [36] | 99.3 [57] | 99.14 [53] | | 68.58 [03] |
| **AM4** | 68.82 [04] | 69.26 [04] | n.a.* | 66.46 [04] | 68.5 [03] | |

*n.a., denotes that no value was given due to low identity and alignment.

**Supplementary Table 3**. Protologues of *Candidatus* Sulfomarinibacter kjeldsenii and *Candidatus* Polarisedimenticola svalbardensis.

| Species name | *Candidatus Sulfomarinibacter kjeldsenii* | *Candidatus Polarisedimenticola svalbardensis* |
|---|---|---|
| Genus etymology | *Sulfomarinibacter* gen. nov. (Sul.fo.ma.ri.ni.bac'ter. L. n. *sulfur* , sulfur; L. adj. *marinus* , marine; L. masc. n. *bacter* (from Gr. n. *bakterion* ), rod; N.L. masc. n. *Sulfomarinibacter* , a sulfur-metabolizing marine rod) | *Polarisedimenticola* gen. nov. (Po.la.ri.se.di.men.ti'co.la. L. adj. *polaris* , polar; L. n. sedimentum, sediment; L. suff. *-cola* , inhabitant, dweller; N.L. masc. n. *Polarisedimenticola* , an inhabitant of polar sediments) |
| Species etymology | *S. kjeldsenii* sp. nov. (kjeld.se'ni.i. N.L. gen. n. *kjeldsenii* , in honour of Kasper Urup Kjeldsen, for his work on the ecology and evolution of sulfur-cycling microorganisms in marine sediments) | *P. svalbardensis* sp. nov. (sval.bar.den'sis. N.L. adj. *svalbardensis* , pertaining to Svalbard) |
| Designation of the type MAG | AM3-C | AM4 |
| MAG accession number | JACXWC000000000 | JACXWD000000000 |
| Genome status[1] | Draft | Draft |
| Estimated genome size | 4.3 Mbp | 3.9 Mbp |
| GC mol% | 60.9 | 62.2 |
| Country of origin | Norway | Norway |
| Region of origin | Svalbard | Svalbard |
| Source of sample | Marine sediment | Marine sediment |
| Sampling date | July, 2016 | July, 2016 |
| Geographic location | Smeerenburgfjorden | Van Keulenfjorden |
| Latitude | 79° 42.83N | 77°32.260′ N |
| Longitude | 11° 05.10E | 15°39.434′ E |
| Water depth | 211 m | 55 m |
| Sediment depth | 5-15 cm | 0-5 cm |
| Sample temperature | − 1.7°C and + 1 to + 3°C, | − 1.7°C and + 1 to + 3°C, |
| Putative energy metabolism | Predicted ability to use cellulose, protein, cyanophycin, hydrogen and acetate. Possible ability to respire nitrous oxide, metal-oxides, tetrathionate, sulfur and sulfite/sulfate, or sulfur disproportionation. | Predicted ability to use protein. Predicted ability to respire oxygen, nitrate, sulfur/polysulfide and metal-oxides |
| Putative relation to oxygen | Anaerobe | Facultative or aerotolerant anaerobe |
| Cell shape | Thin rods, ~2 x 0.5 microns, visualized by CARD-FISH. | So far unidentified. |

[1]See Table 1 for further genome information.

**Supp. Table 4.** Key gene/protein annotations. (all RAST files with protein encoding genes (pegs) corresponding to the numbers in this table can be found in Figshare: https://doi.org/10.6084/m9.figshare.13022675.v1)

| Pathway/function | Gene | AM1 | AM2 | AM3-A | AM3-B | AM3-C | AM4 | . | dsr-harbouring ThM_scaffold_807 | dsr-harbouring scaffold_276 |
|---|---|---|---|---|---|---|---|---|---|---|
| | | | | | | | | . | | |
| **Dissimilatory sulfate reduction (sulfate<->sulfide )** | *dsrA* | . | . | . | 1691-2 | . | . | . | 1 | . |
| | *dsrB* | . | 961 | . | 1693 | 1570 | . | . | 2 | 3234 |
| | *dsrC* | . | 196 | . | 1055 | 17 | . | . | 9 | 3227 |
| | *dsrD* | . | . | . | 1694 | . | . | . | 3 | 3233 |
| | *dsrL* | . | . | . | 1059 | . | . | . | 5 | 3231 |
| | *dsrN* | . | . | . | 1058 | 20 | . | . | 6 | 3230 |
| | *dsrM* | . | 198 | . | 1053 | 1478 | . | . | 11 | 3225 |
| | *dsrK* | 1947? | 1020 | . | . | 1875 | . | . | 12 | 3224 |
| | *dsrJ* | . | . | . | . | 1138 | . | . | 13 | 3223 |
| | *dsrO* | . | . | . | . | . | . | . | 14 | 3222 |
| | *dsrP* | 1931 | . | . | . | . | . | . | 15 | 3221 |
| | | | | | | | | . | | |
| | sulfate adenylyltransferase (*sat* ) | 1835 | . | . | 1231 | 1489 | . | . | | |
| | adenylylsulfate reductase (*aprA* ) | 1842 | . | . | 1633 | 588 | . | . | | |
| | adenylylsulfate reductase (*aprB* ) | 1459 | . | . | 1634 | 589 | . | . | | |
| | | | | | | | | . | | |
| | quinone-interacting membrane oxidoreductase complex *(qmoA)* | 1839 | 415 | . | 1630 | 2274 | . | . | | |
| | quinone-interacting membrane oxidoreductase complex (*qmoB* ) | 1838 | 416 | . | 1629 | 2273 | . | . | | |
| | quinone-interacting membrane oxidoreductase complex *(qmoC)* | 1836 | 417 | . | . | . | . | . | | |
| | | | | | | | | . | | |
| **Sulfur/thiosulfate reduction** | polysulfide/thiosulfate reductase (*psrA* ) | . | . | . | . | . | 1867 | . | | |
| | polysulfide/thiosulfate reductase (*psrB* ) | . | . | . | . | . | 1868 | . | | |
| | polysulfide/thiosulfate reductase (*psrC* ) | . | . | . | . | . | 1869 | . | | |
| | | | | | | | | . | | |
| **Sulfur/polysulfide reduction-hydrogenase** | sulfyhdrogenase, alpha subunit ([NiFe] hydrogenase subunit alpha) | . | . | . | . | . | 1009 | . | | |
| | sulfyhdrogenase, beta subunit (anaerobic sulfite reductase subunit A) | . | . | . | . | . | 1013 | . | | |
| | sulfyhdrogenase, gamma subunit (anaerobic sulfite reductase subunit B) | . | . | . | . | . | 1011 | . | | |
| | sulfyhdrogenase, delta subunit (hydrogenase/oxidoreductase) | . | . | . | . | . | 1010 | . | | |
| | | | | | | | | . | | |
| **Tetrathionate reduction** | cytochrome tetrathionate reductase (*otrA* ) | 699/700 | . | . | . | 1334 | . | . | | |
| | | | | | | | | . | | |

| | | | | | | | | |
|---|---|---|---|---|---|---|---|---|
| **Complex-Iron-Sulfur-Molybdoenzymes (CISM)** | tetrathionate reductase subunit A (*ttrA* ) | . | . | 1080 | . | . | . | . |
| | tetrathionate reductase subunit B (*ttrB* ) | . | . | 1081 | . | . | . | . |
| | | | | | | | | . |
| | CISM protein with unknown function | . | . | . | 235 | . | 362, 2746 | . |
| | arsenate reductase, subunit A (*arrA* ) | . | . | . | . | . | 2793 | . |
| | arsenate reductase, subunit B (*arrB* ) | . | . | . | . | . | 2792 | . |
| | arsenate reductase, subunit C (*arrC* ) | . | . | . | . | . | 2791 | . |
| | | | | | | | | . |
| | cytochrome c oxidase aa3, subunit II | . | 328 | . | . | 1373 | 1316 | . |
| **Cytochromes (oxygen reduction)** | cytochrome c oxidase aa3, subunit I | 1384 | 987 | . | . | . | 1317 | . |
| | cytochrome c oxidase aa3, subunit III | 1385 | 986 | . | 56 | 1449 | 1318 | . |
| | cytochrome c oxidase aa3, subunit III (copy 2) | . | . | . | . | 1451 | 1320 | . |
| | cytochrome c oxidase aa3, subunit IV | 1386 | . | . | . | 1450 | 319 | . |
| | | | | | | | | . |
| | cytochrome c oxidase cbb3, subunit I | . | 1192 | . | 2613 | 832 | 1323 | . |
| | cytochrome c oxidase cbb3, subunit II | . | 1192 | . | 2613 | 832 | 1323 | . |
| | cytochrome c oxidase cbb3, subunit IV | . | 1193 | . | 2614 | 831 | 1324 | . |
| | cytochrome c oxidase cbb3, subunit III | . | 1194 | . | 2615 | 830 | 1325 | . |
| | | | | | | | | . |
| **Extracellular cytochromes & associated genes** | multi-heme cytochrome c | 893 | . | . | . | 3164 | 1123 | . |
| | NHL repeat domain protein (β-barrel) | 894 | . | . | . | 3165 | 1124 | . |
| | extracellular cytochrome c | 895 | . | . | . | 3166 | 1125 | . |
| | peptidyl-prolyl cis-trans isomerase-like (β-barrel chaperon) | 896 | . | . | . | 3167 | . | . |
| | hypothetical protein (β-barrel) | . | . | . | . | 3168 | 1127 | . |
| | hypothetical protein (β-barrel) | . | . | . | . | 3169 | . | . |
| | multi-heme cytochrome c | . | . | . | . | 3170 | 1128 | . |
| | multi-heme cytochrome c | | | | | | 1129 | . |
| | | | | | | | | . |
| **OmcS-like cytochromes and associated** | cytochrome c family protein (*omcS* -like) | . | . | . | . | . | 2398 | . |
| | hypothetical protein | . | . | . | . | . | 2399 | . |
| | lipoprotein, putative | . | . | . | . | . | 2400 | . |
| | hypothetical protein | . | . | . | . | . | 2401 | . |
| | foldase protein PrsA precursor (EC 5.2.1.8) | . | . | . | . | . | 2402 | . |
| | cytochrome c family protein | . | . | . | . | . | 2403 | . |
| | cytochrome c family protein | . | . | . | . | . | 2404 | . |
| | NHL repeat protein (β-barrel) | . | . | . | . | . | 2405 | . |
| | NHL repeat domain protein (β-barrel) | . | . | . | . | . | 2406 | . |
| | Cytochrome c family protein (*omcS* -like) | . | . | . | . | . | 2407 | . |
| | | | | | | | | . |

| | | | | | | | | |
|---|---|---|---|---|---|---|---|---|
| | nitrous oxide reductase (*nosZ* , type-II) | 148 | . | . | 69 | 1027 | . | . |
| **Reduction of nitrogenous compunds** | probable copper chaperone (*nosL* ) | 149 | . | . | 68 | . | . | . |
| | nitrous oxide reductase maturation protein (*nosD* ) | 1754 | 857 | . | 66 | 80 | . | . |
| | nitrous oxide reductase maturation transmembrane protein (*nosY* ) | 1751 | 854 | . | 63 | . | . | . |
| | | | | | | | | . |
| | nitrate reductase, periplasmic catalytic subunit (*napA* ) | . | . | . | . | . | 258 | . |
| | nitrate reductase cytochrome c-type subunit (*napB* ) | . | . | . | . | . | 257 | . |
| | nitrate reductase, ferredoxin-type protein (*napF* ) | . | . | . | . | . | 256 | . |
| | | | | | | | | . |
| **Reductive dehalogenation** | reductive dehalogenase homologous protein A, catalytic subunit (*rdhA* ) | 42 | . | . | . | 1850 | . | . |
| | reductive dehalogenase homologous protein B, membrane-anchor (*rdhB* ) | . | . | . | . | 1851 | . | . |
| | | | | | | | | . |
| **Electron flows & energy conservation** | heterodisulfide reductase, subunit A | 293 | . | 391 | 2210 | 2020 | . | . |
| | heterodisulfide reductase, subunit B | 294 | . | 392 | 2209 | 2021 | . | . |
| | heterodisulfide reductase, subunit C | . | . | 393 | 2208 | 2022 | . | . |
| | (additional Hdr subunits are encoded adjacent to above genes) | | | | | | | . |
| | | | | | | | | . |
| | high-molecular-weight cytochrome c3 | 1948 | . | . | . | . | . | . |
| | | | | | | | | . |
| | NADH ubiquinone oxidoreductase chain A | 1526 | 866 | . | 1374 | 987 | 2103 | . |
| | NADH-ubiquinone oxidoreductase chain B | 1987 | 865 | . | 2142 | 1628 | 2104 | . |
| | NADH-ubiquinone oxidoreductase chain C | 1989 | . | . | 2141 | 1627 | 2105 | . |
| | NADH-ubiquinone oxidoreductase chain D | 1990-1 | . | . | 2140 | 1866 | 2106 | . |
| | NADH-ubiquinone oxidoreductase chain H | 1993 | 549 | . | 2139 | 624 | 2107 | . |
| | NADH-ubiquinone oxidoreductase chain I | 1994 | 449 | . | 2138 | 625 | 2108 | . |
| | NADH-ubiquinone oxidoreductase chain J | 1995 | 448 | . | 2137 | 626 | 2109 | . |
| | NADH-ubiquinone oxidoreductase chain K | 1996 | 447 | . | 2136 | 627 | 2110 | . |
| | NADH-ubiquinone oxidoreductase chain L | 1425 | 1456 | 1357 | 2135 | 628 | 2111 | . |
| | NADH-ubiquinone oxidoreductase chain M | 13 | 1455 | . | 2134 | 629 | 2112 | . |
| | NADH-ubiquinone oxidoreductase chain N | 578 | 1116 | 1356 | 2133 | 630 | 2113 | . |
| | NADH-ubiquinone oxidoreductase chain E | 1525 | 1099 | . | 1371 | 986 | . | . |
| | NADH-ubiquinone oxidoreductase chain F | 1524 | 1314 | . | 1372 | 985 | 92 | . |
| | NADH-ubiquinone oxidoreductase chain G | 1523 | 1315 | . | 1373 | 984 | . | . |
| | | | | | | | | . |
| | Na(+)-translocating NADH-quinone reductase subunit A | . | . | 278 | 2930 | 118 | 2698 | . |
| | Na(+)-translocating NADH-quinone reductase subunit A | 1069 | 1159 | 277 | 2931 | 117 | 2699 | . |

| | | | | | | | | |
|---|---|---|---|---|---|---|---|---|
| | Na(+)-translocating NADH-quinone reductase subunit B | 1070 | 1158 | 276 | 2932 | 116 | . | . |
| | Na(+)-translocating NADH-quinone reductase subunit C | . | . | 275 | 2933 | 115 | . | . |
| | Na(+)-translocating NADH-quinone reductase subunit D | . | . | 274 | 2934 | 114 | . | . |
| | Na(+)-translocating NADH-quinone reductase subunit E | . | . | 273 | 2935 | 113 | . | . |
| | Na(+)-translocating NADH-quinone reductase subunit F | 1726 | . | 272 | 2936 | 112 | 2245 | . |
| | | | | | | | | . |
| **Fermentation** | acetate kinase | . | . | . | 34 | 755 | 1646 | . |
| | phosphate acetyltransferase | 437 | . | . | 1978 | 309 | 2062 | . |
| | | | | | | | | . |
| **Carbohydrate degradation/modifications** | cellulase A | . | 628 | 250 | 1313 | 2253 | . | . |
| | cellobiose phosphorylase | . | . | . | . | 2593 | . | . |
| | cellobiose phosphotransferase system YdjC-like protein | . | . | 1569 | 1214 | . | . | . |
| | | | | | | | | . |
| | Carbohydrate active enzymes (glycoside hydrolases) | 338 | 1336 | 1570 | 1213 | 157 | 1241 | . |
| | | 756 | 1354 | 251 | 2853 | 2341 | 1795 | . |
| | | 156 | 628 | 1572 | 264 | 2255 | 1954 | . |
| | | 1363 | 771 | 840 | 1099 | 15 | 1967 | . |
| | | 669 | 705 | 906 | 729 | 2252 | 2311 | . |
| | | 336 | 143 | 456 | 273 | 1441 | 3120 | . |
| | | 1362 | 282 | 332 | 2915 | 1209 | 3121 | . |
| | | 747 | . | 1050 | 1211 | 1226 | 3122 | . |
| | | 788 | . | 1056 | 1315 | 1715 | 347 | . |
| | | 484 | . | 1022 | 859 | 802 | 918 | . |
| | | 743 | . | 571 | 1663 | 476 | . | . |
| | | 367 | . | 1085 | 1507 | 257 | . | . |
| | | 70 | . | . | 1134 | 2054 | . | . |
| | | . | . | . | 2682 | 1474 | . | . |
| | | . | . | . | 1475 | 1591 | . | . |
| | | . | . | . | 3114 | 2444 | . | . |
| | | . | . | . | 1351 | 2302 | . | . |
| | | . | . | . | 2404 | . | . | . |
| | | . | . | . | 238 | . | . | . |
| | | | | | | | | . |
| **Cyanophycin degradation** | cyanophycinase | . | . | . | 1887 | 1417 | . | . |
| | isoaspartyl dipeptidase | . | . | . | . | 698 | . | . |
| | arginine deiminase | 201 | . | 517 | 2247 | 747 | 2134 | . |
| | ornithine carbamoyltransferase | 2634 | 324 | 1337 | 824 | 2614 | 2489 | . |

| | | | | | | | | |
|---|---|---|---|---|---|---|---|---|
| | carbamate kinase | 1139 | . | 521 | 2252 | 1944 | 2352 | . |
| | aspartate ammonia-lyase (catabolic) | . | . | . | 1678 | 2840 | . | . |
| | | | | | | | | . |
| **Hydrogen and formate utilisation/convertions** | NAD-dependent formate dehydrogenase alpha subunit | . | 950 | . | . | 2772 | 90 | . |
| | formate dehydrogenase beta subunit | . | 948 | . | . | 2770 | 92 | . |
| | Pyruvate-formate lyase | . | 803 | 1346 | . | 1666 | . | . |
| | Pyruvate-formate lyase, activating enzyme | 431 | . | 1345 | 1926 | 1665 | . | . |
| | [Ni/Fe] up-take hydrogenase, large subunit | 1602 | 1083 | . | 1930 | 2955 | 2448 | . |
| | [Ni/Fe] up-take hydrogenase, small subunit | 1601 | 1084 | . | . | 2956 | 2449 | . |
| | [Ni/Fe] up-take hydrogenase, cytochrome b subunit | . | 1084 | . | 1928 | 2957 | 2447 | . |
| | | | | | | | | . |
| | lysyl endopeptidase (EC 3.4.21.50) | . | . | . | . | 1383 | 307 | . |
| **Protein degradation, catabolic predicted (extracellular/periplasmic)** | peptidase with M28 domain | . | . | . | . | . | 83 | . |
| | alkaline serine protease | . | . | . | . | . | 1753 | . |
| | cold-active serine alkaline protease | . | . | . | . | . | 2259 | . |
| | serine protease, subtilase family | . | . | . | . | . | 2586 | . |
| | serine protease | . | . | . | . | . | 405 | . |
| | Peptidases_S8_Kp43_protease | . | . | . | . | . | 306 | . |
| | Peptidases_S8_Subtilisin_like | . | . | . | . | . | 2391 | . |
| | extracellular alkaline serine protease | . | . | . | . | 2367 | . | . |
| | Zn-dependent dipeptidase | . | . | . | . | 2368 | . | . |
| | beta-lactamase (D-aminoacylase) | . | . | . | . | 2369 | . | . |

**Supplementary Table 5**. Best 25 BLASTP hits from NCBI RefSeq database to OmcS-like proteins from MAG AM4.

**MAG AM4, peg.2398**

| Best hit | Max Score | Total Score | Query Coverage | e- value | % identity | Accession no. |
|---|---|---|---|---|---|---|
| hypothetical protein [candidate division Zixibacteria bacterium] | 389 | 389 | 99% | 7.00E-132 | 59.49% | WP_156049117.1 |
| hypothetical protein [candidate division Zixibacteria bacterium] | 382 | 382 | 97% | 4.00E-129 | 54.25% | WP_156049015.1 |
| hypothetical protein [Geopsychrobacter electrodiphilus] | 187 | 187 | 95% | 4.00E-52 | 34.96% | WP_020675668.1 |
| hypothetical protein [Geobacter daltonii] | 182 | 182 | 97% | 4.00E-50 | 32.87% | WP_012648237.1 |
| cytochrome C [Geomonas oryzae] | 177 | 177 | 97% | 9.00E-48 | 32.22% | WP_129126951.1 |
| cytochrome C [Geomonas terrae] | 176 | 176 | 97% | 2.00E-47 | 32.22% | WP_135871229.1 |
| hypothetical protein [Geomonas edaphica] | 174 | 174 | 97% | 4.00E-47 | 33.33% | WP_136513827.1 |
| cytochrome C [Desulfuromonas sp. AOP6] | 173 | 173 | 96% | 4.00E-47 | 35.53% | WP_155876307.1 |
| cytochrome C [Geomonas ferrireducens] | 174 | 174 | 97% | 6.00E-47 | 32.00% | WP_136525908.1 |
| cytochrome C [Geomonas edaphica] | 174 | 174 | 97% | 7.00E-47 | 32.00% | WP_136513815.1 |
| hypothetical protein [Pelobacter seleniigenes] | 172 | 172 | 97% | 1.00E-46 | 33.50% | WP_051689551.1 |
| cytochrome C [Desulfuromonas acetexigens] | 171 | 171 | 97% | 7.00E-46 | 32.84% | WP_140396604.1 |
| hypothetical protein [Geobacter uraniireducens] | 170 | 170 | 92% | 2.00E-45 | 31.94% | WP_011938882.1 |
| hypothetical protein [Geomonas oryzae] | 169 | 169 | 97% | 2.00E-45 | 32.61% | WP_129126940.1 |
| c-type cytochrome OmcS [Geobacter anodireducens] | 170 | 170 | 97% | 2.00E-45 | 32.57% | WP_066358290.1 |
| cytochrome C [Geobacter anodireducens] | 169 | 169 | 92% | 3.00E-45 | 32.93% | WP_066357145.1 |
| cytochrome c [Geobacter soli] | 169 | 169 | 92% | 4.00E-45 | 32.93% | WP_039643446.1 |
| c-type cytochrome OmcS [Geobacter soli] | 169 | 169 | 97% | 4.00E-45 | 32.56% | WP_039648128.1 |
| hypothetical protein [Geomonas terrae] | 169 | 169 | 97% | 4.00E-45 | 32.45% | WP_135871207.1 |
| c-type cytochrome OmcS [Geobacter sulfurreducens] | 169 | 169 | 97% | 5.00E-45 | 32.72% | WP_010943141.1 |
| hypothetical protein [Geopsychrobacter electrodiphilus] | 167 | 167 | 92% | 5.00E-45 | 34.29% | WP_020675667.1 |
| cytochrome C [Geomonas ferrireducens] | 167 | 167 | 98% | 1.00E-44 | 33.87% | WP_136525916.1 |
| hypothetical protein [Geomonas ferrireducens] | 167 | 167 | 97% | 2.00E-44 | 31.96% | WP_136525917.1 |
| cytochrome C [Geomonas oryzae] | 167 | 167 | 97% | 2.00E-44 | 33.49% | WP_129126941.1 |
| c-type cytochrome OmcS [Geobacter sulfurreducens] | 167 | 167 | 97% | 2.00E-44 | 32.26% | WP_119334099.1 |

**MAG AM4, peg.2407**

| Best hit | Max Score | Total Score | Query Coverage | e- value | % identity | Accession no. |
|---|---|---|---|---|---|---|
| hypothetical protein [candidate division Zixibacteria bacterium] | 268 | 268 | 99% | 2.00E-84 | 46.11% | WP_156049015.1 |
| hypothetical protein [candidate division Zixibacteria bacterium] | 230 | 230 | 98% | 1.00E-69 | 42.36% | WP_156049117.1 |
| cytochrome C [Desulfuromonas sp. AOP6] | 181 | 181 | 100% | 3.00E-50 | 35.13% | WP_155876307.1 |
| hypothetical protein [Geopsychrobacter electrodiphilus] | 173 | 173 | 99% | 5.00E-47 | 35.22% | WP_020675668.1 |
| cytochrome C [Desulfuromonas acetexigens] | 165 | 165 | 95% | 9.00E-44 | 36.46% | WP_140396604.1 |
| cytochrome C [Geoalkalibacter subterraneus] | 155 | 155 | 97% | 2.00E-40 | 33.51% | WP_144401977.1 |
| hypothetical protein [Geoalkalibacter subterraneus] | 153 | 153 | 98% | 8.00E-40 | 35.80% | WP_158414055.1 |
| cytochrome C [Pelobacter seleniigenes] | 150 | 150 | 87% | 9.00E-39 | 36.63% | WP_155005876.1 |
| hypothetical protein [Malonomonas rubra] | 150 | 150 | 97% | 2.00E-38 | 34.18% | WP_072908292.1 |
| cytochrome c [Geobacter sulfurreducens] | 149 | 149 | 96% | 7.00E-38 | 33.85% | WP_010941362.1 |
| cytochrome C [Geobacter sp. DSM 9736] | 149 | 149 | 95% | 8.00E-38 | 32.37% | WP_088536830.1 |
| hypothetical protein [Geobacter daltonii] | 148 | 148 | 97% | 3.00E-37 | 29.74% | WP_012648237.1 |
| hypothetical protein [Desulfuromonas acetexigens] | 145 | 145 | 98% | 1.00E-36 | 31.94% | WP_092055201.1 |
| hypothetical protein [Geomonas ferrireducens] | 145 | 145 | 98% | 2.00E-36 | 29.88% | WP_136525917.1 |
| hypothetical protein [Geobacter sulfurreducens] | 145 | 145 | 96% | 2.00E-36 | 33.68% | WP_014551388.1 |
| cytochrome C [Geobacter bremensis] | 145 | 145 | 90% | 2.00E-36 | 33.33% | WP_156912219.1 |
| hypothetical protein [Anaeromyxobacter dehalogenans] | 144 | 144 | 83% | 5.00E-36 | 33.63% | WP_015934404.1 |
| hypothetical protein [Anaeromyxobacter dehalogenans] | 144 | 144 | 88% | 6.00E-36 | 32.96% | WP_011422115.1 |
| hypothetical protein [Geobacter uraniireducens] | 144 | 144 | 97% | 8.00E-36 | 32.17% | WP_011938897.1 |
| hypothetical protein [Geomonas edaphica] | 143 | 143 | 98% | 9.00E-36 | 29.70% | WP_136513827.1 |
| cytochrome c [Geobacter bremensis] | 144 | 144 | 97% | 1.00E-35 | 31.68% | WP_051292969.1 |
| hypothetical protein [Geobacter daltonii] | 143 | 143 | 94% | 1.00E-35 | 31.31% | WP_012648226.1 |
| hypothetical protein [Geomonas oryzae] | 142 | 142 | 92% | 3.00E-35 | 31.15% | WP_129126940.1 |
| hypothetical protein [Anaeromyxobacter sp. PSR-1] | 142 | 142 | 88% | 4.00E-35 | 32.68% | WP_059436742.1 |
| cytochrome C [Geobacter bemidjiensis] | 141 | 141 | 95% | 7.00E-35 | 33.82% | WP_148212949.1 |

**Supplementary Table 6.** Gylcoside hydrolases detected by dbCAN2 analyses.

| | MAG | Gylcoside hydrolases (dbCAN) | Protein encoding genes | %GHs |
|---|---|---|---|---|
| **Marine** | AM1 | 13 | 2134 | 0.61% |
| (this study) | AM2 | 7 | 1482 | 0.47% |
| | AM3-A | 12 | 1596 | 0.75% |
| | AM3-B | 20 | 3197 | 0.63% |
| | AM3-C | 18 | 3255 | 0.55% |
| | AM4 | 17 | 3174 | 0.54% |
| **Terrestrial** | SbA1 | 82 | 5235 | 1.57% |
| **(*dsr* -harbouring)** | SbA3 | 169 | 9407 | 1.80% |
| (Hausmann *et al* . 2018) | SbA4 | 127 | 10701 | 1.19% |
| | SbA5 | 147 | 5114 | 2.87% |
| | SbA6 | 73 | 3650 | 2.00% |
| | SbA7 | 16 | 3230 | 0.50% |

**Supplementary Table 7.** Percentages of metagenomic read pairs mapped to MAGs (>99% identity). Values are factored for MAG completeness estimates. SM, Smeerenburgfjorden. VK, Van Keulenfjorden.

| | SM, St. J, 04 cmbsf | SM, St. J, 10-15 cmbsf | SM, St. J, 17 cmbsf | VK, St. AC 18 cmbsf |
|---|---|---|---|---|
| **AM1** | 0.013% | 0.062% | 0.170% | 0.002% |
| **AM2** | 0.023% | 0.137% | 0.148% | 0.004% |
| **AM3-A** | 0.011% | 0.078% | 0.196% | 0.011% |
| **AM3-B** | 0.018% | 0.118% | 0.268% | 0.016% |
| **AM3-C** | 0.018% | 0.122% | 0.279% | 0.017% |
| **AM4** | 0.000% | 0.000% | 0.000% | 0.215% |
