## Supplementary Figures 1-18 for "Novel taxa of Acidobacteriota involved in seafloor sulfur cycling"

A.

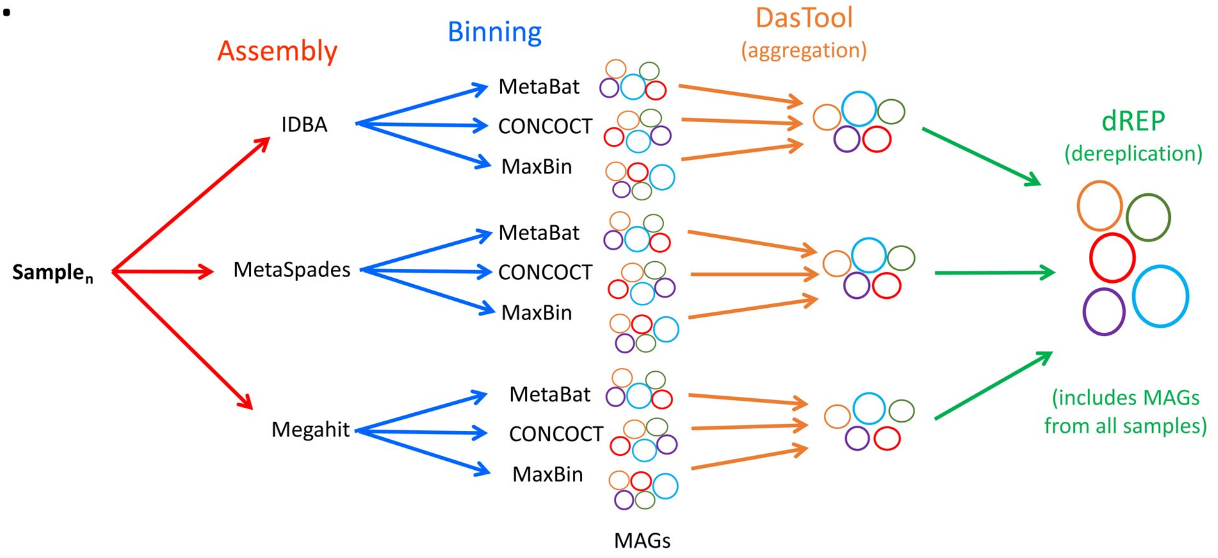


B.

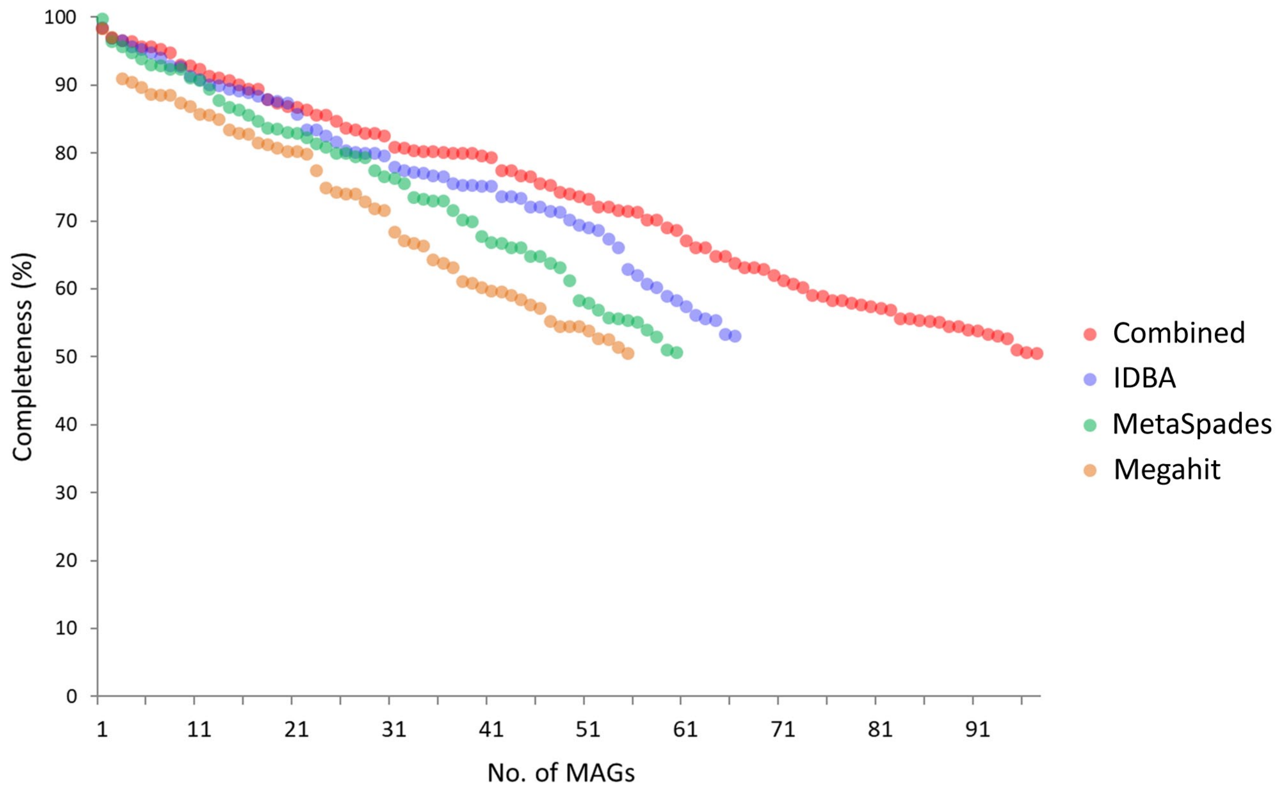


**Supplementary Figure 1**. A) Outline of the metagenomic binning strategy. B) Plot of completeness of MAGs (CheckM). Comparisons are derived from binning from single assemblies (IDBA or MetaSpades or Megahit), versus binning from multiple assemblies of each sample (outlined in panel A).

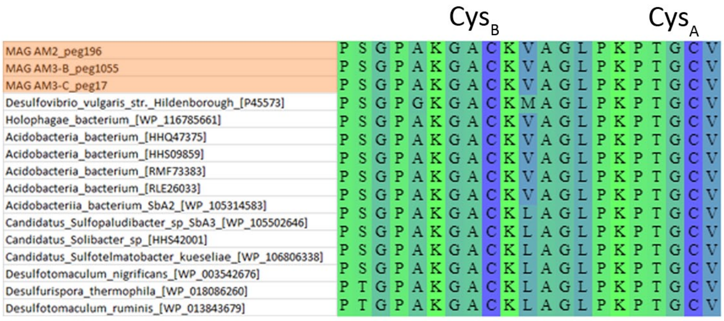


**Supplementary. Fig. 2.** Alignment of dissimilatory DsrC cysteine motifs. Sub-section (C-terminus) of alignment of DsrC proteins, showing two conserved cysteine residues (dark purple) that are present in dissimilatory versions of the enzymes.

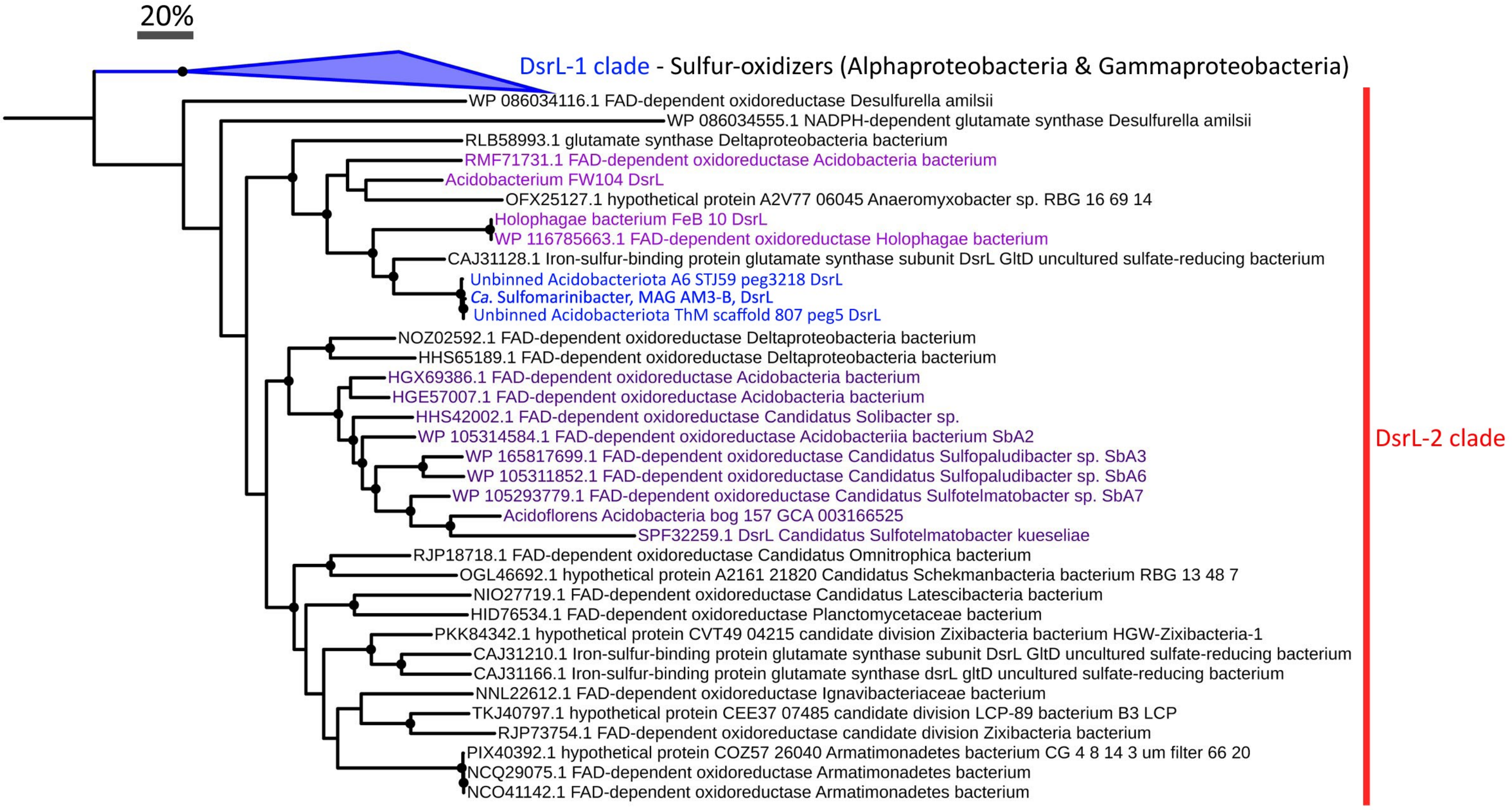


**Supplementary Figure 3A.** Phylogenetic tree of DsrL proteins. The sequences from MAGs recovered in this study are highlighted in blue. Other Acidobacteriota DsrL are highlighted in purple. Bootstrap values >50% are presented on nodes as black-filled circles. The scale bar represents 20% sequence divergence.

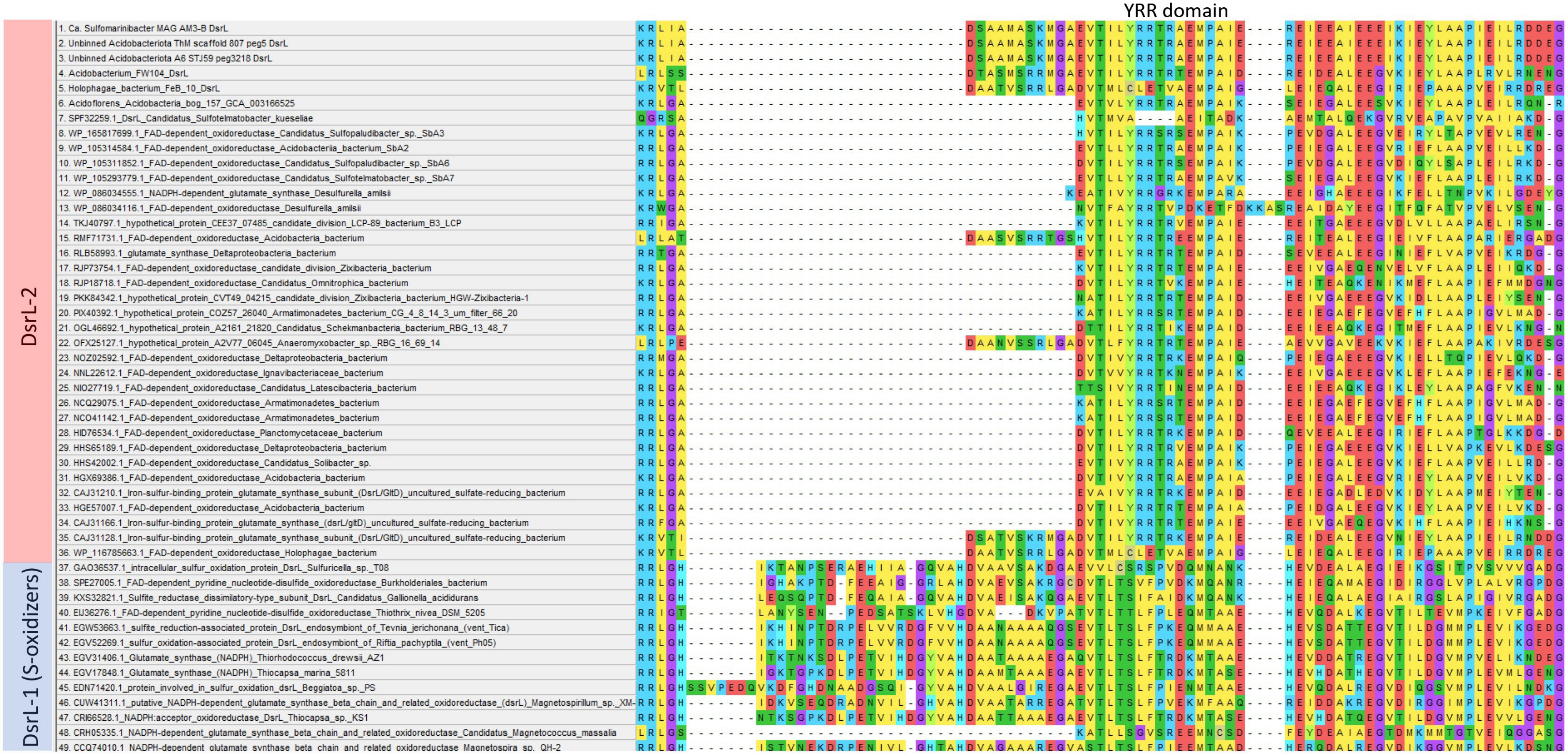

**Supplementary Figure 3B)**. Alignment of DsrL proteins. A subsection of the whole DsrL alignnment is shown to highlight YRR amino acids of putative NAD(P)H-binding domain.

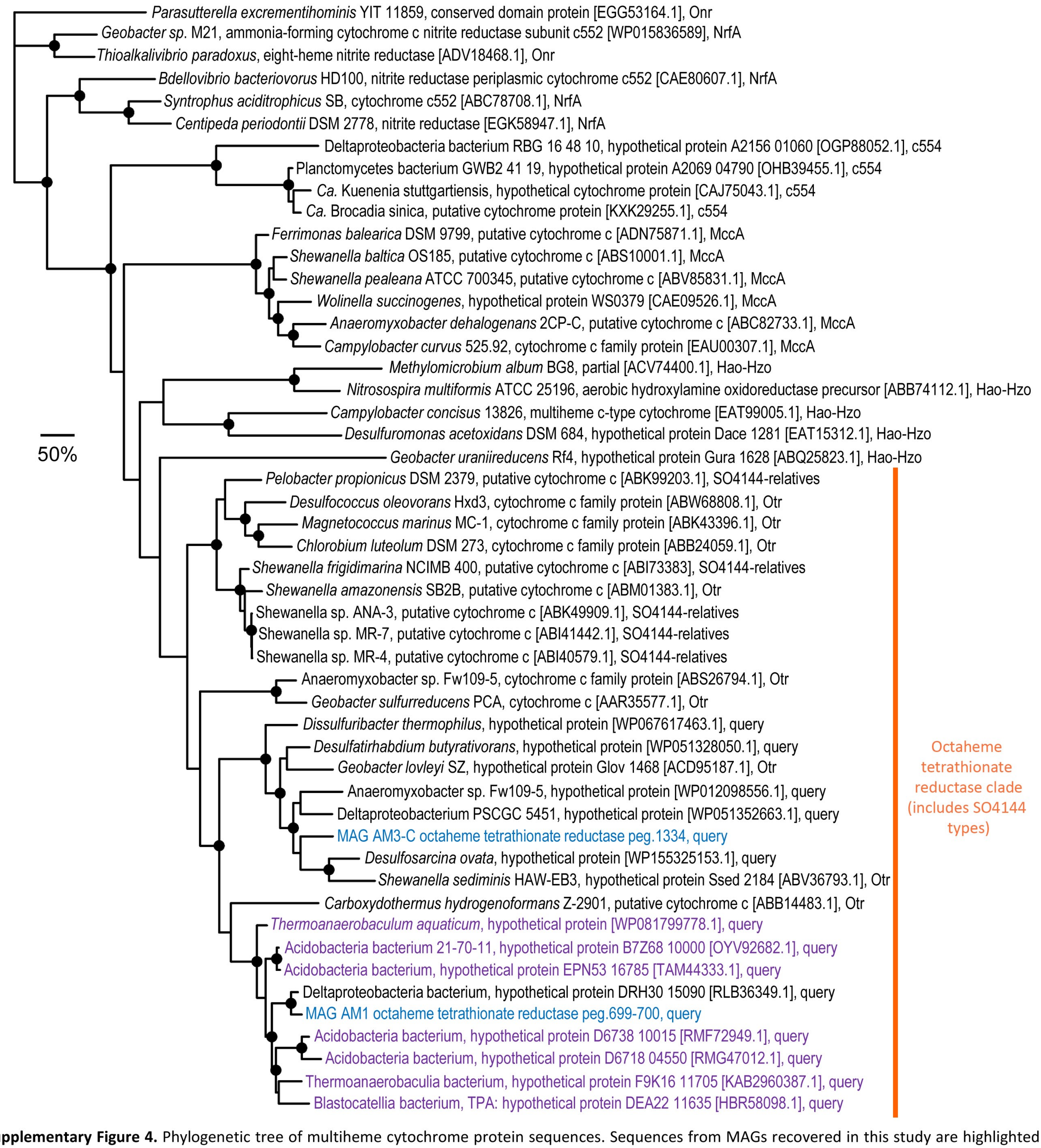


**Supplementary Figure 4.** Phylogenetic tree of multiheme cytochrome protein sequences. Sequences from MAGs recovered in this study are highlighted in blue. Sequences from other Acidobacteriota are highlighted in purple. Reference sequences were retrieved from Kern et al. 2011, and from best BLASTP hits to our MAG-derived sequences. Functional assignments are labelled at the end of each leaf label. Bootstrap values with >90% are indicated with filled black circles on nodes. The scale bar represents 50% sequence divergence. Genbank accessions are presented in parentheses. NrfA = respiratory cytochrome c nitrite reductase, Onr = octaheme cytochrome c nitrite reductase, Hao/Hzo = octahaem hydroxylamine oxidoreductase/hydrazine oxidoreductase, MccA = cytochrome c sulfite reductase, and Otr = octaheme tetrathionate reductase.

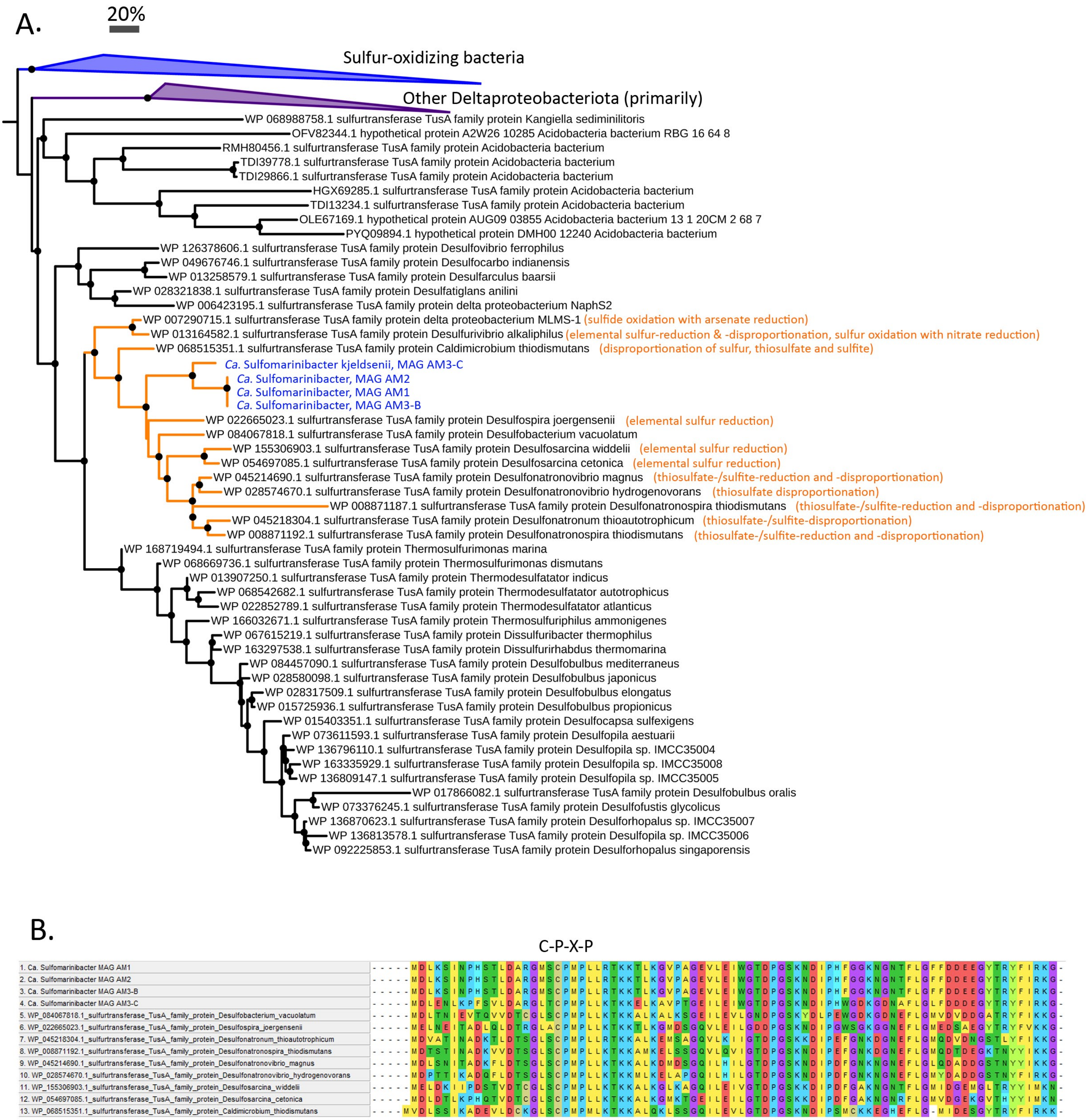


**Supplementary Figure 5. A)** Phylogenetic tree of TusA proteins. The sequences from MAGs recovered in this study are highlighted in blue. The orange branch indicates TusA proteins from anaerobic organisms known to reduce or disproportionate sulfur cycle intermediates and that had TusA related to the Aciobacteriota TusA. Descriptions of sulfur metabolisms related to reduction or disproportionation of sulfur cycle intermediates are presented in parenthesis for TusA related to TusA from MAGs recovered in this study. Bootstrap values >50% are presented on nodes as black-filled circles. The scale bar represents 20% sequence divergence. **B)** Alignment of TusA proteins from marine Acidobacteriota showing Cys-Pro-X-Pro sulfane sulfur-binding domains.

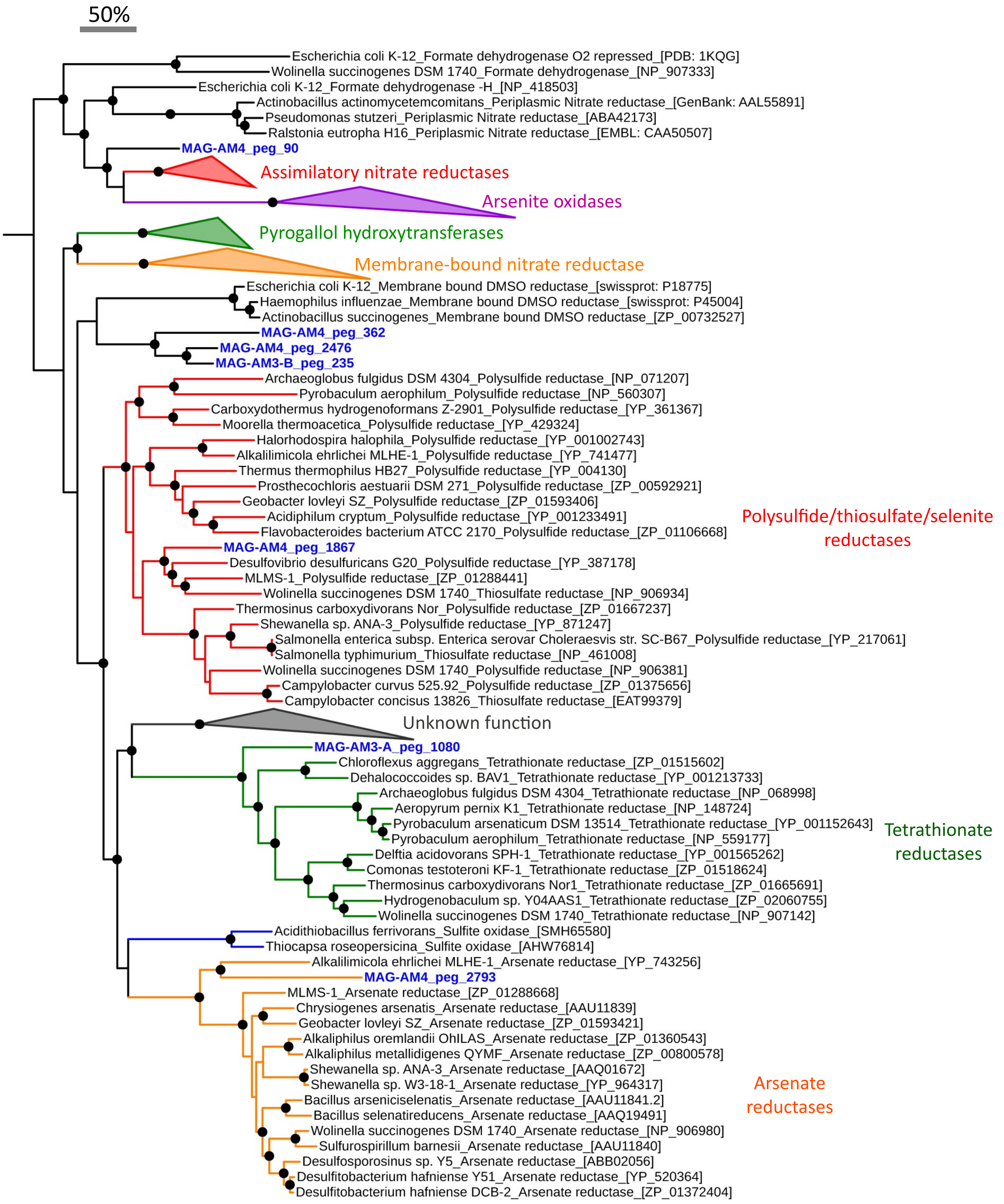


**Supplementary Figure 6.** Phylogenetic tree of complex iron–sulfur molybdoenzyme (CISM) family proteins. The sequences from the MAGs recovered in this study are highlighted in blue. Reference sequences were obtained from Duval et. al. 2008, as well as selected additional sequences. Bootstrap values >90% are presented on nodes as black-filled circles. The scale bar represents 50% sequence divergence.

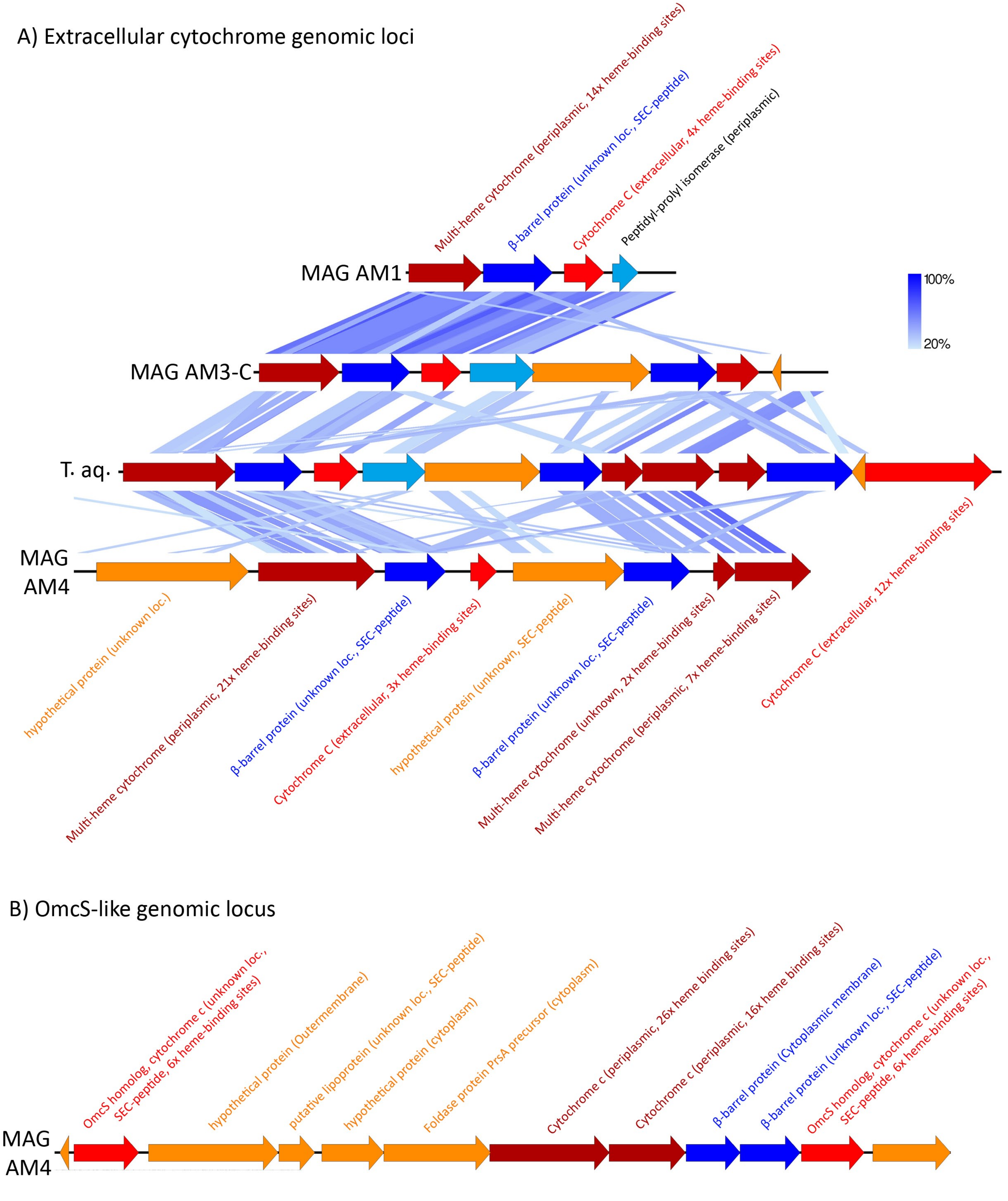


**Supplementary Figure 7**. **A)** Schematic of gene organisation and synteny of extracellular cytochrome-rich genomic loci among Acidobacteria MAGs (AM1, AM3-C and AM4) and *Thermoanaerobaculum aquaticum* (T. aq.). **B)** Schematic of gene organisation of genomic loci encoding OmcS-like proteins in MAG AM4. Shaded blue lines indicate degree of sequence similarity as determined by tblastx within EasyFig (Sullivan et al., 2011). Subcellular location predictions and number of heme-binding sites (CXXCH) are indicated in parentheses. SEC-peptides for Sec secretion systems were searched in proteins with 'unknown' location predictions using PRED-TAT (Bagos et al., 2011).
.

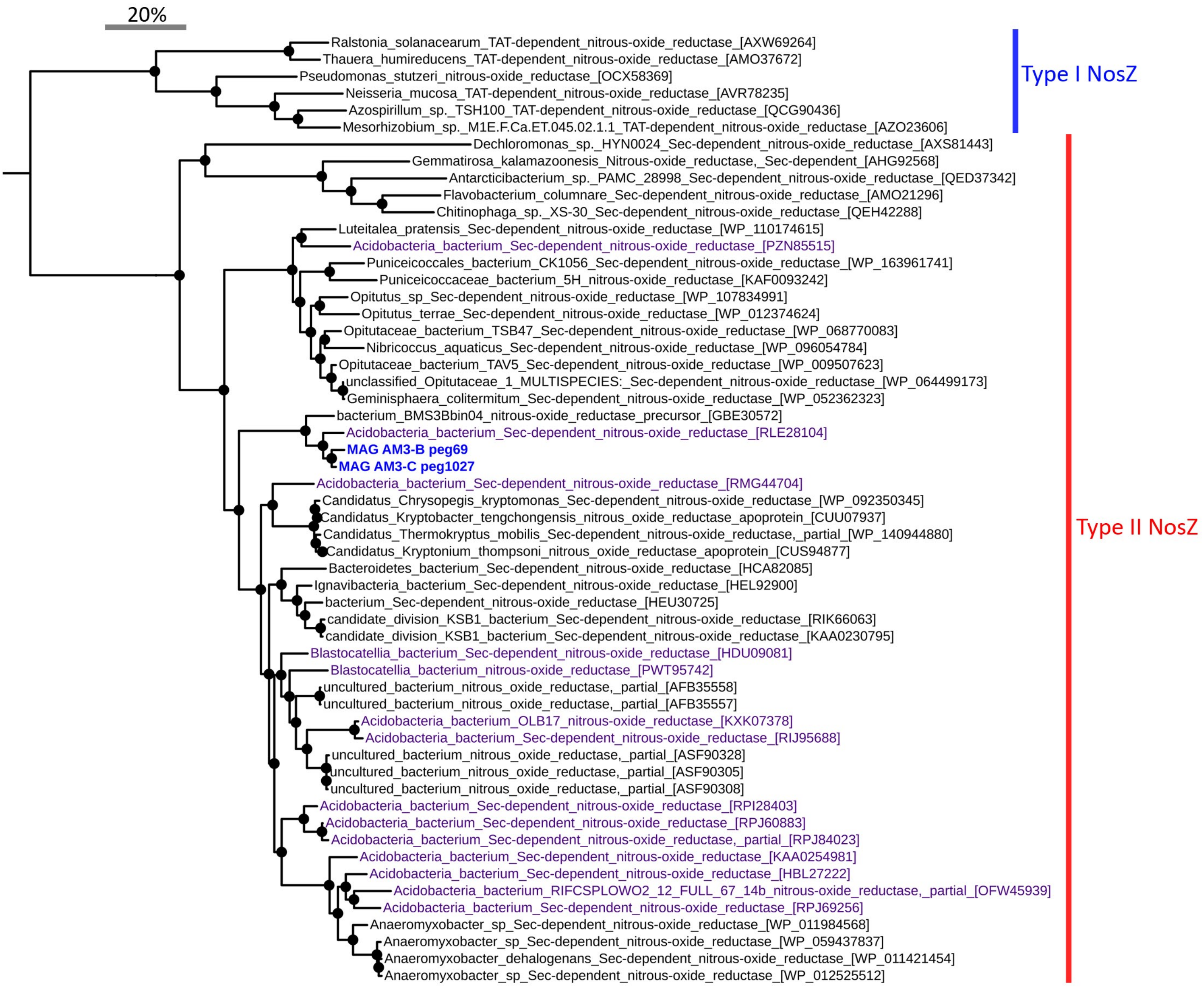


**Supplementary Figure 8.** Phylogenetic tree of nitrous oxide reductases (NosZ). Sequences from MAGs recovered in this study are highlighted in blue. The NosZ from MAG AM1 was omitted due to short sequence length, although it was most similar to the NosZ from AM3-B and AM3-C (>90% amino acid identity from 190 amino acids). Sequences from other Acidobacteriota are highlighted in purple. Clade of 'type I NosZ' = blue, and clade of 'type II NosZ' = red. Reference sequences were retrieved from the top 50 best BLASTP hits to the NosZ from MAG AM3-C were included. Genbank accessions are presented in parentheses. Black circles on nodes represent bootstraps values >90%. The scale bar represents 20% sequence divergence.

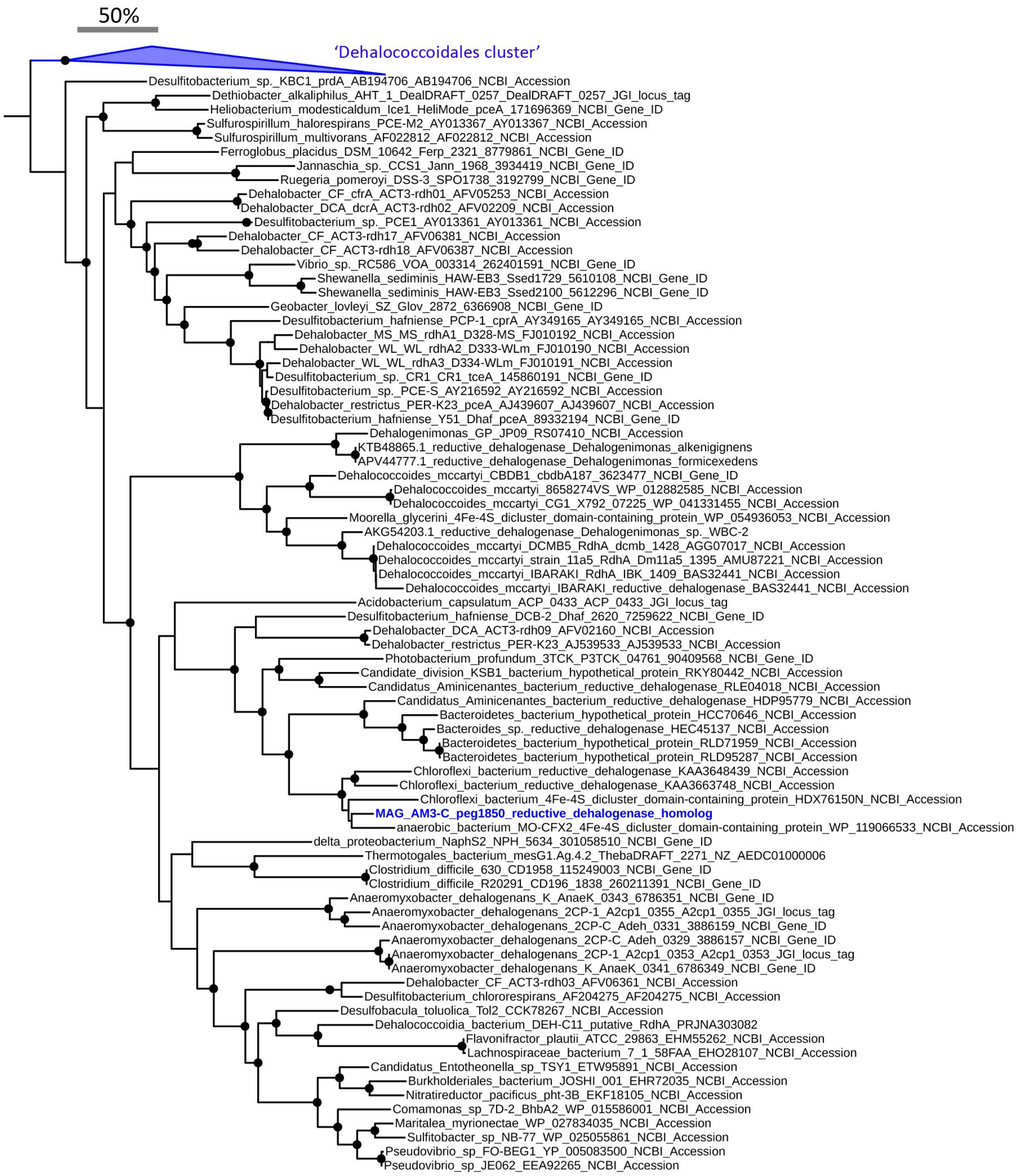


**Supplementary Figure 9.** Phylogenetic tree of reductive dehalogenase homolog A (RdhA) proteins. The sequence from the MAG recovered in this study are highlighted in blue. Reference sequences were obtained from Hug et. al. 2013, and the top 10 best BLASTP hits to the RdhA from MAG AM3-C were also included. The RdhA sequence of MAG AM1 was not included due to the truncated protein sequence, although it was most similar to the RdhA from MAG AM3-C (>87% amino acid identity from 120 amino acids). Black circles on nodes represent bootstraps values >90%. The scale bar represents 50% sequence divergence.

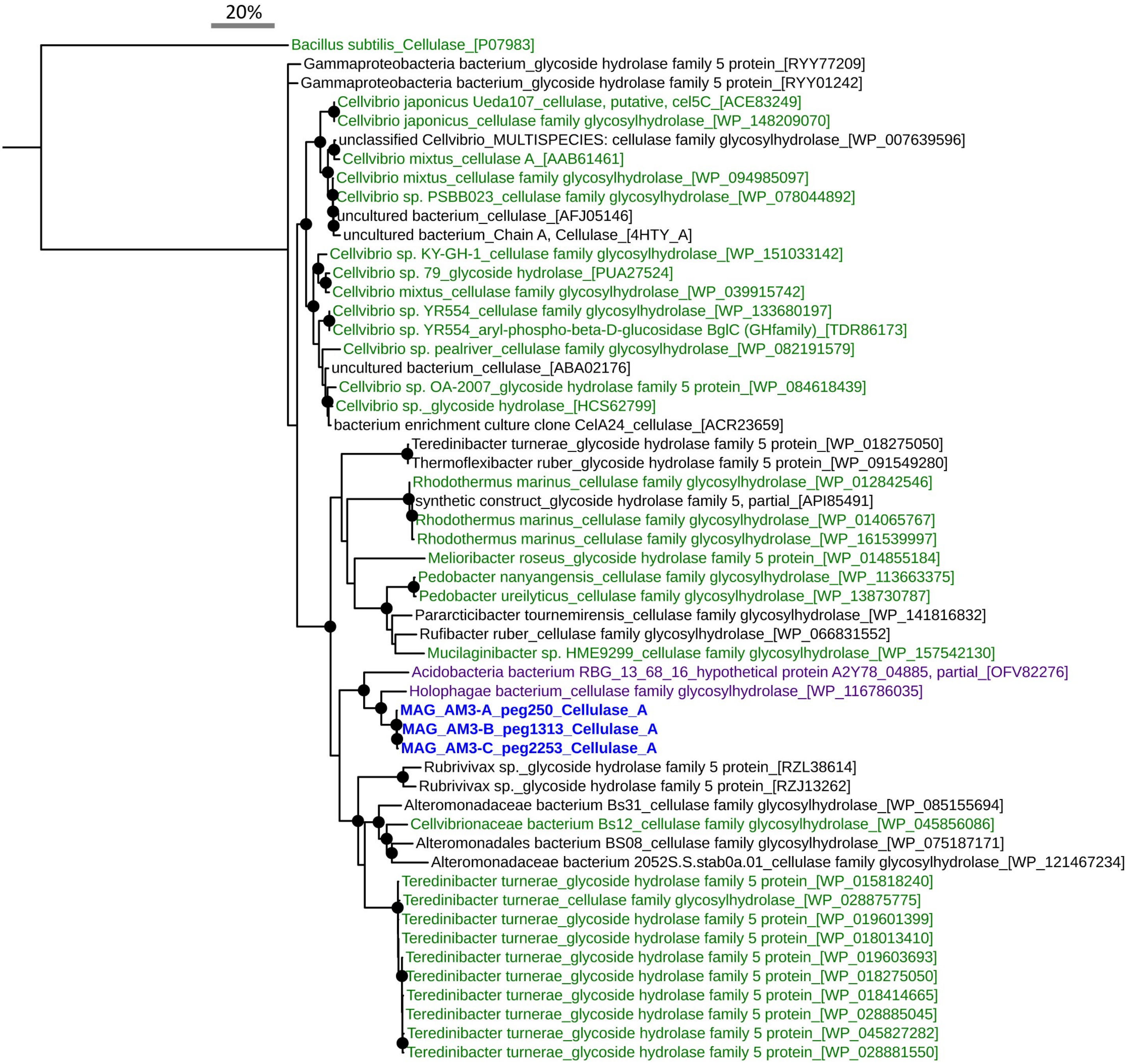


**Supplementary Figure 10.** Phylogenetic tree of cellulase A-like proteins. Sequences from MAGs recovered in this study are highlighted in blue. Sequences from other Acidobacteria are highlighted in purple. Sequences from genera or species known to perform cellulose degradation are highlighted in green. Reference sequences were retrieved from the top 50 best BLASTP hits to the cellulase A from MAG AM3-C. Genbank accessions are presented in parentheses. The tree was rooted with the cellulase A of *Bacillus subtilis*. Black circles on nodes represent bootstraps values >90%. The scale bar represents 20% sequence divergence.

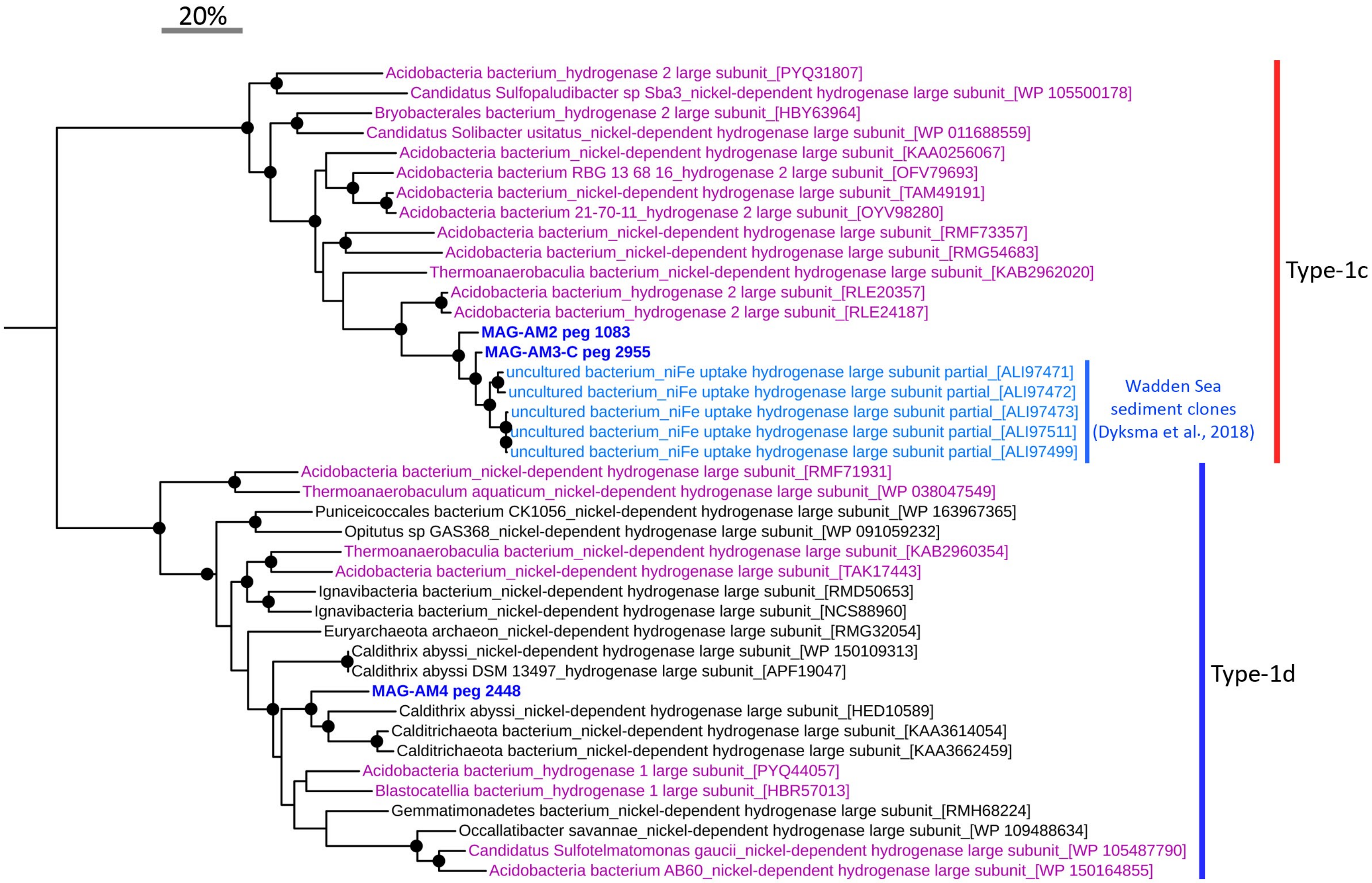


**Supplementary Figure 11.** Phylogenetic tree of [NiFe]-hydrogenase large subunit proteins. The sequences from the MAGs recovered in this study are highlighted in dark blue. Sequences from other Acidobacteriota are highlighted in purple. Sequences from PCR-derived amplicons from tidal flat sediments (Dyksma et al., 2018) are highlighted in light blue. Reference sequences were derived from best BLASTP hits from NCBI-nr database. Hydrogenase 'types' were determined using HydDB (Søndergaard et al., 2016). Black circles on nodes represent bootstraps values >90%. The scale bar represents 20% sequence divergence.

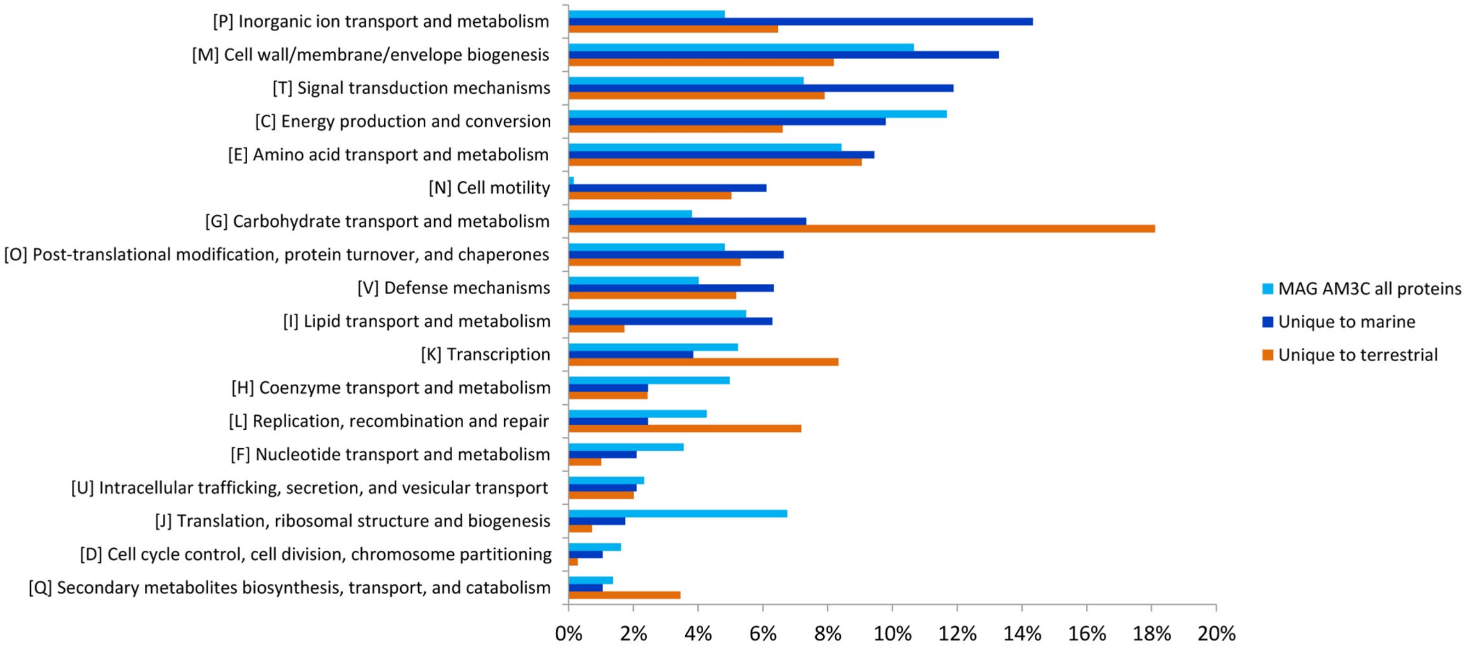


**Supplementary Figure 12.** Comparisons of COG classifications of proteins representing unique ortholog groups (OGs) from marine versus terrestrial *dsr*-harbouring Acidobacteriota. OGs unique to each group of genomes were determined using OrthoFinder (Emms and Kelly 2019). Proteins were compared from the six MAGs recovered in this study, versus proteins from the seven MAGs recovered by Hausmann *et al*. 2018. Letters in parenthesis represent standard COG codes.

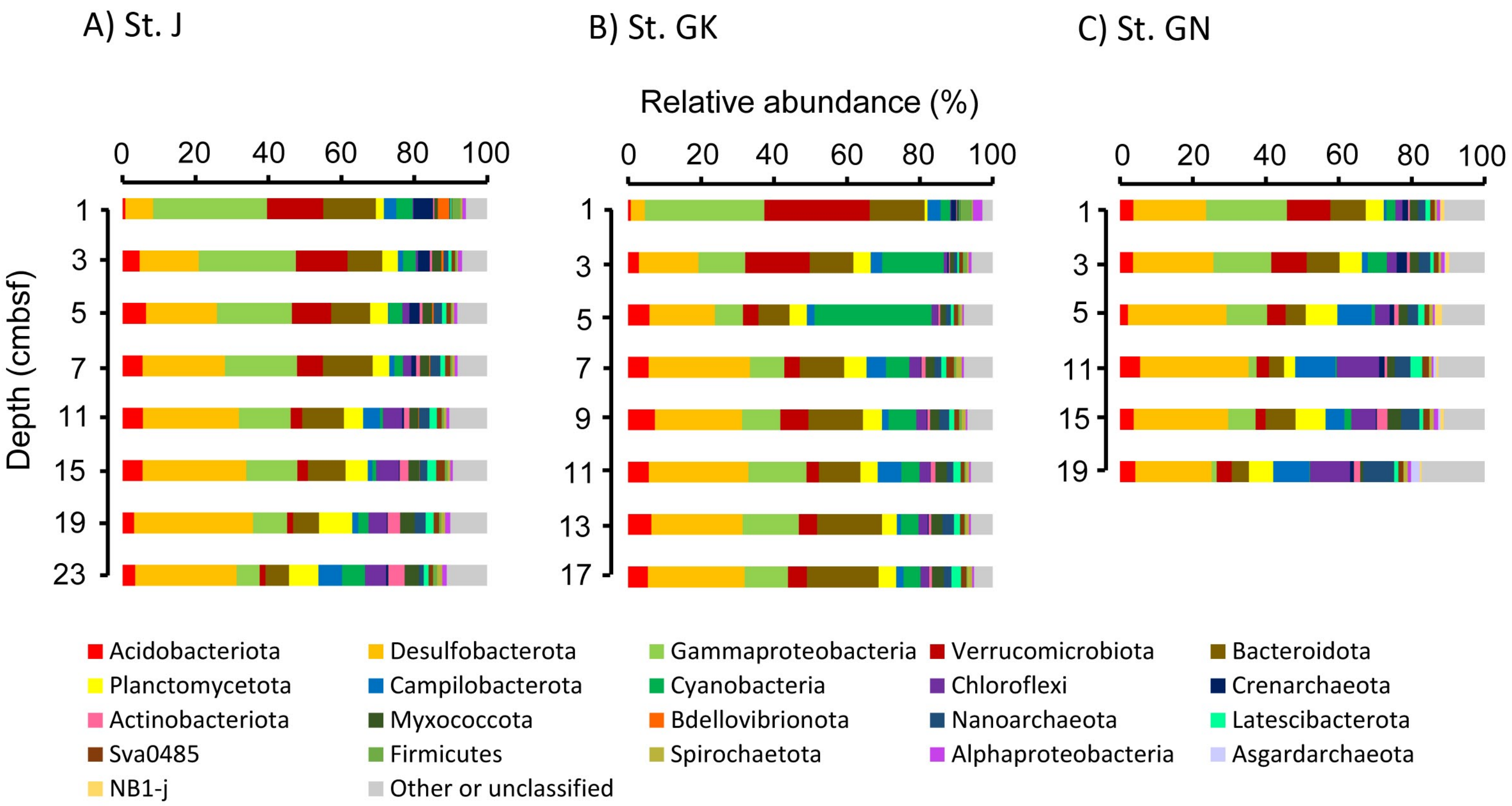


**Supplementary Figure 13.** Microbial community composition of Smeerenburgfjorden sediments. Relative abundance of 16S rRNA ASVs is derived from amplicon sequencing of the 16S rRNA gene. Taxa that are less abundant than 1% or are unclassified are shown in grey. Depth is shown in centimeters below seafloor (cmbsf) for A) Station J, B) Station GK, and C) Station GN.

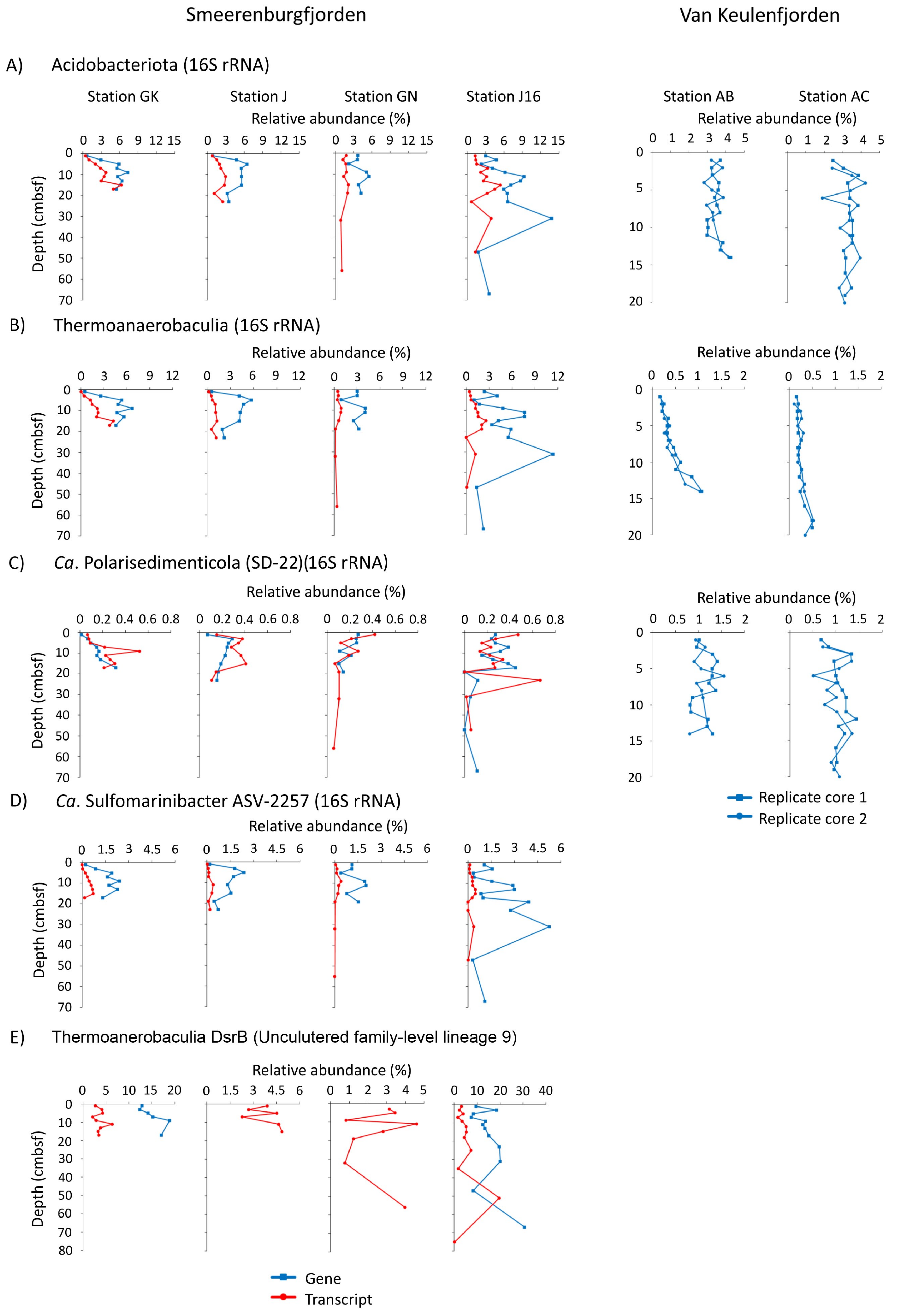


**Supplementary Figure 14.** Relative abundances of 16S rRNA and DsrB genes and transcripts for Svalbard sediments. A) Acidobacteriota, B) Thermoanaerobaculia, C) *Ca.* Polarisedimenticola (SD-22), D) *Ca.* Sulfomarinibacter ASV-2257, E) Thermoanaerobaculia DsrB (uncultured family-level lineage 9). Smeerenburgfjorden stations GK, J and GN were sampled in June 2017, and J16 was sampled in July 2016. Replicate cores from Van Keulenfjorden stations AB and AC are derived from Buongiorno et al., 2019.

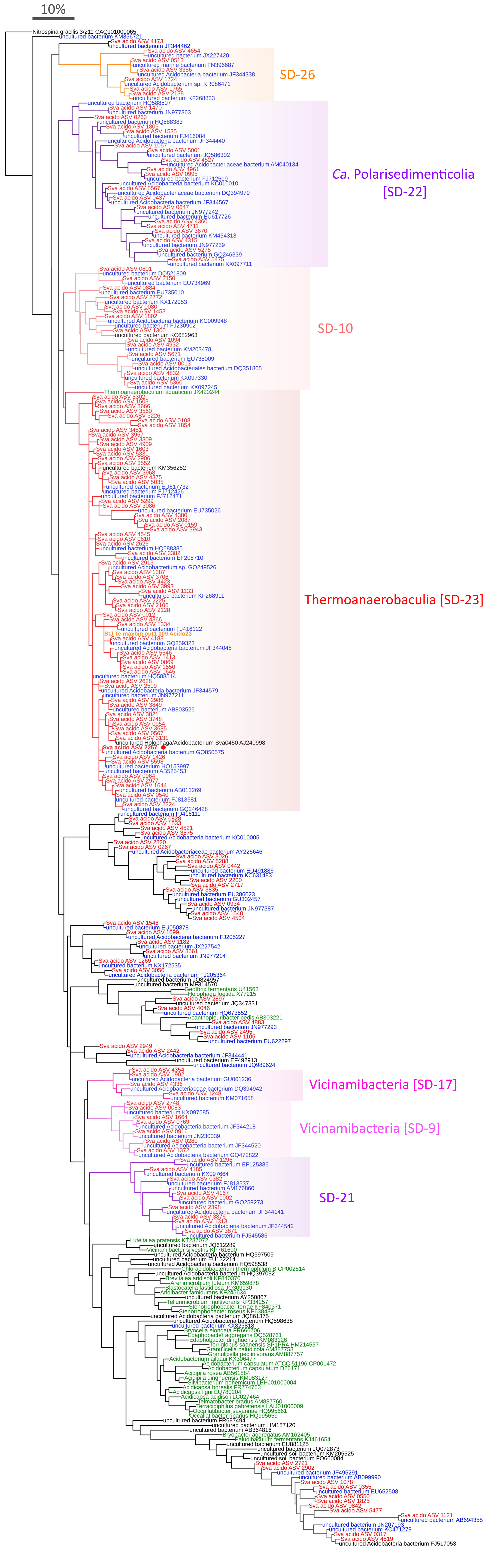


**Supplementary Figure 15.** Phylogenetic tree of 16S rRNA gene sequences. Red leaves are from all acidobacterial amplicon-derived sequences retrieved in this study from Smeerenburgfjorden, Svalbard. The red dot denotes the most abundant ASV in the dataset. The orange leaf represents the 16S rRNA sequence recovered from an acidobacterial metagenome-assembled genome (this study). Blue leaves represent sequences derived from marine environments and present in the SILVA database v138. The blue leaves were selected as the closest relatives of the query sequences. Green leaves represent cultivated Acidobacteria. SD = 'sub-division' as per SILVA database (v138). The tree was built as a consensus of three maximum-likelihood methods (see Materials and Methods). The scale bar represents 10% sequence divergence.

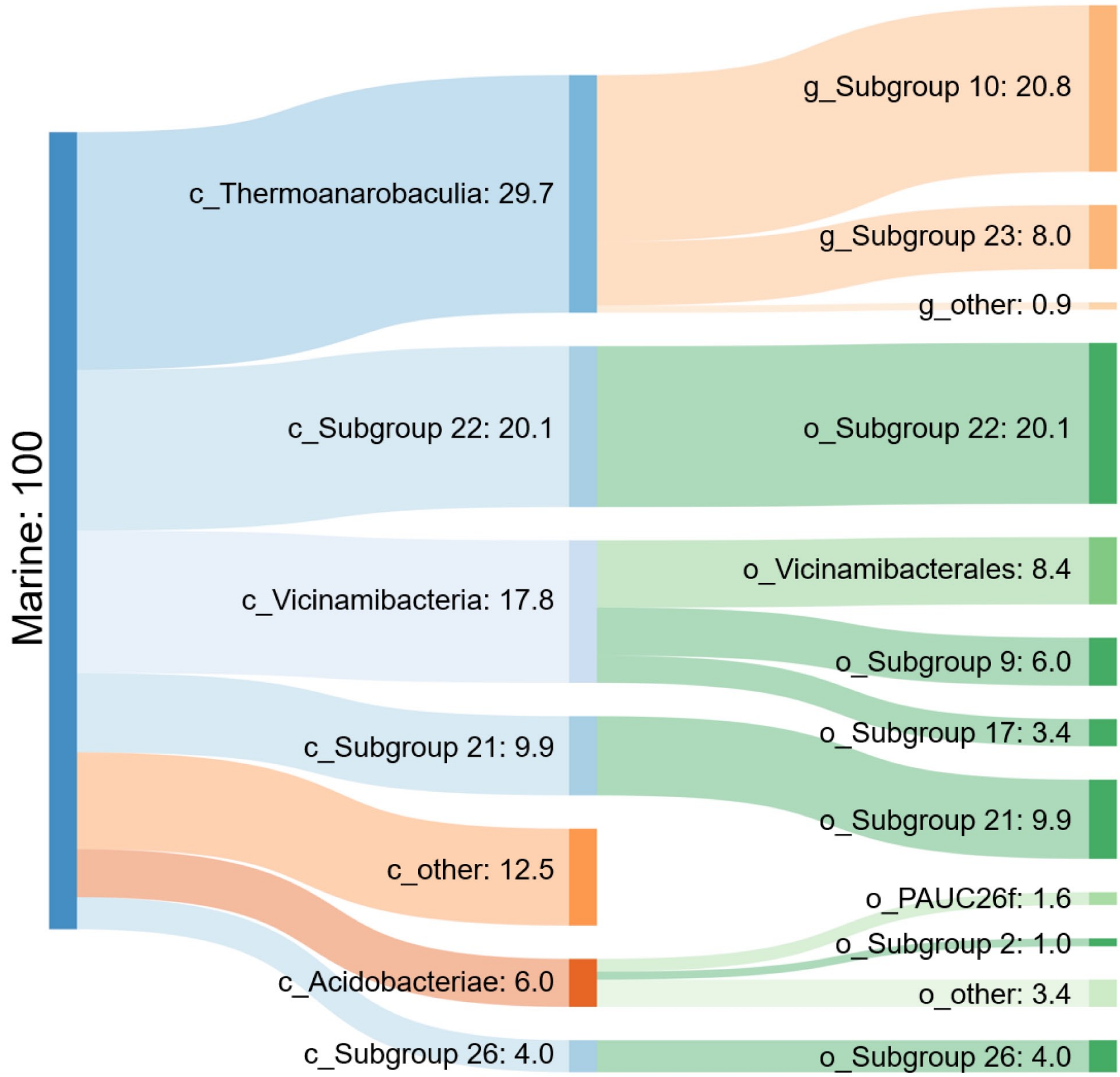


**Supplementary Figure 16.** Sankey diagram of marine Acidobacteriota 16S rRNA gene sequences and their major taxonomic classifications. The SILVA database (v138 NR) was the source of the marine Acidobacteriota 16S rRNA gene sequences (n=771). Numbers refer to the percentage of all marine sediment-derived Acidobacteriota sequences.

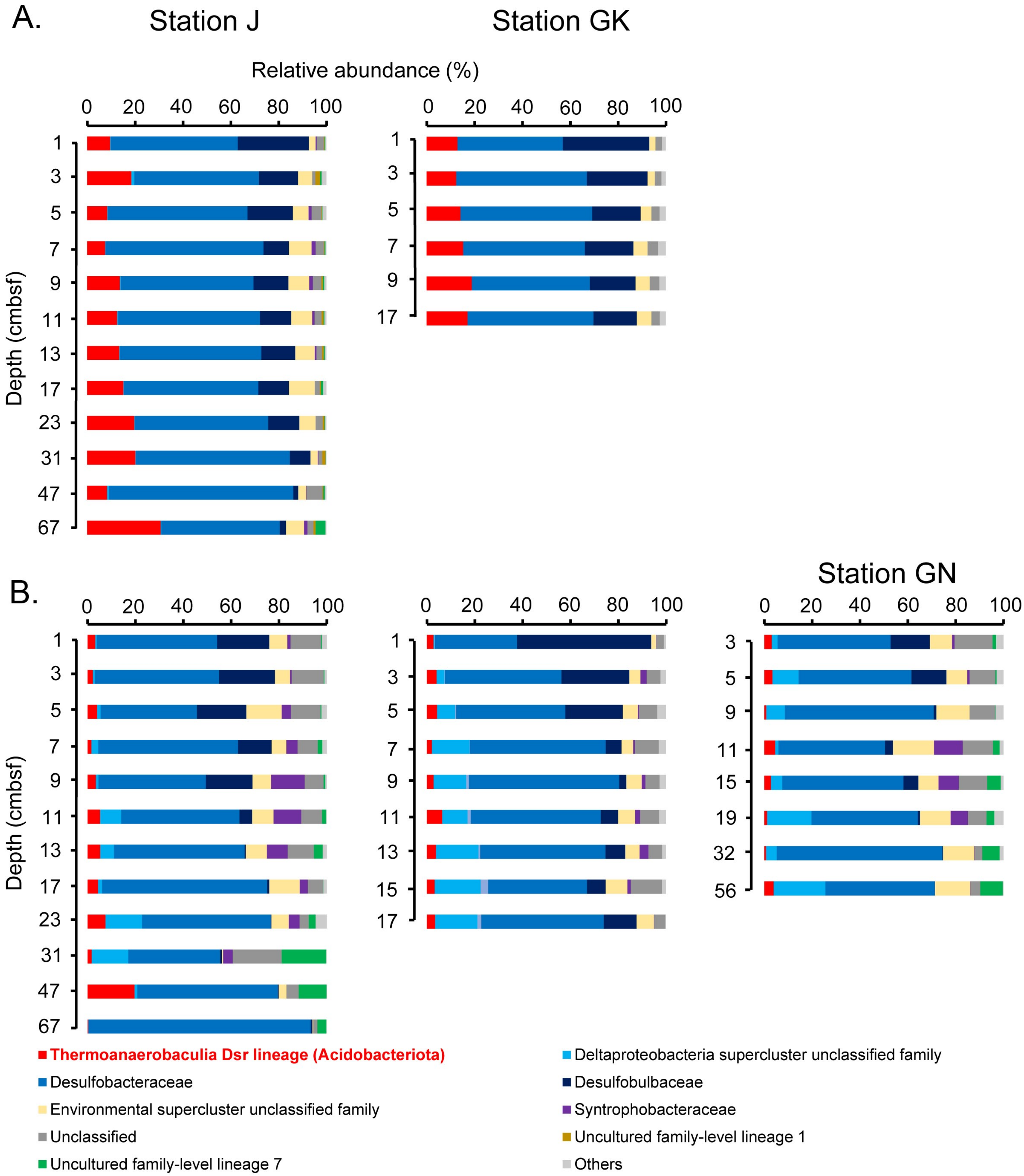


**Supplementary Figure 17**. Community compositions of *dsrB*-harboring microorganisms in Svalbard sediments. A) Compositions determined from *dsrB*-gene (DNA) amplicon sequencing. B) Compositions determined from *dsrB*-transcript (cDNA) amplicon sequencing. Groups that are less abundant than 1% are grouped as ‘Others’.

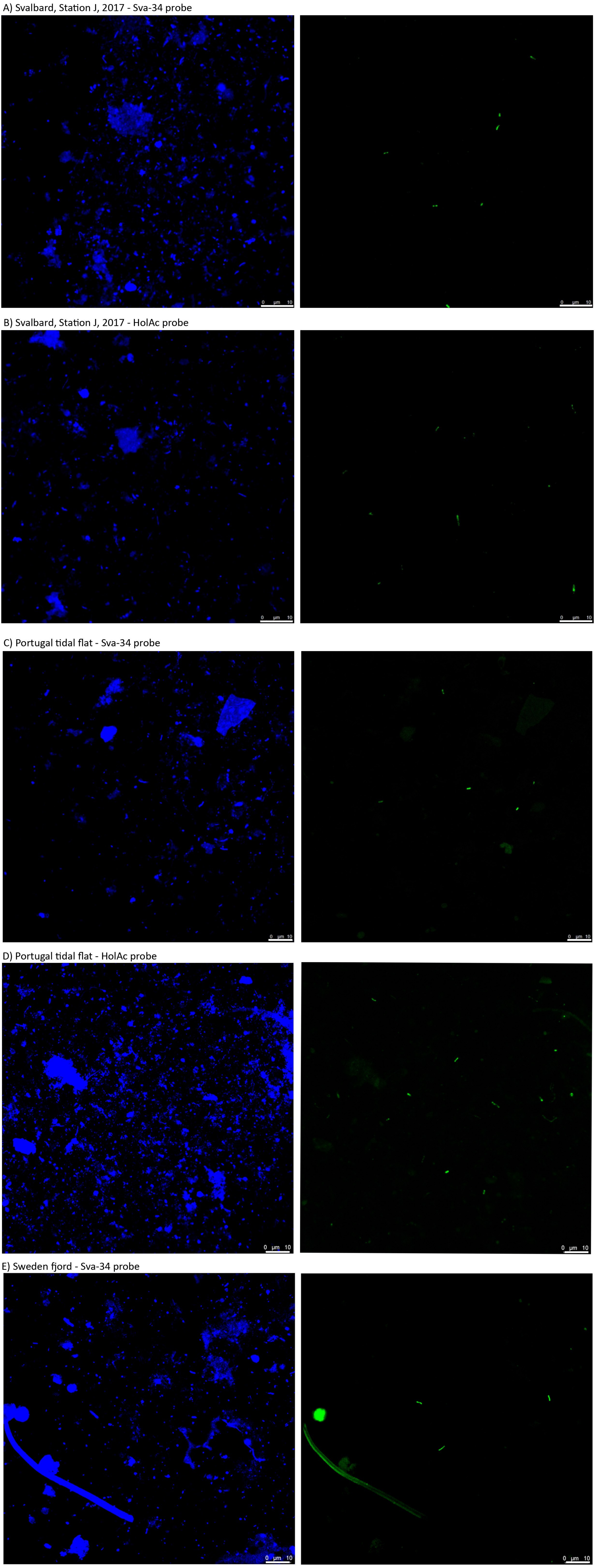


**Supplementary Figure 18.** CARD-FISH images of Acidobacteriota from marine sediments. Sediment locations and probes used are listed above panels A-E. Panels with blue cells are DAPI stained, panels with green cells are CARD-FISH hybridised cells from corresponding fields of view. White scale bars represent 10 µm.
